# DNA Methylation Dynamics of Dose-dependent Acute Exercise, Training Adaptation, and Detraining

**DOI:** 10.1101/2025.04.22.650067

**Authors:** Manoj Hariharan, Sahil Patel, Haili Song, Abid Rehman, Cesar Barragan, Anna Bartlett, Rosa Castanon, Joseph Nery, Vincent Rothenberg, Huaming Chen, Wei Tian, Wubin Ding, Wenliang Wang, Jeremy McAdam, Zachary Graham, Kaleen Lavin, Marcas Bamman, Timothy Broderick, Joseph Ecker

## Abstract

Exercise and diet are direct physical contributors to human health, wellness, resilience, and performance^1–5^. Endurance and resistance training are known to improve healthspan through various biological processes such as mitochondrial function^6–8^, telomere maintenance^9^, and inflammaging^10^. Although several training prescriptions have been defined with specific merits ^1,10–20^, the long-term effects of these in terms of their molecular alterations have not yet been well explored. In this study, we focus on two combined endurance and resistance training programs: (1) traditional moderate-intensity continuous endurance and resistance exercise (TRAD) and (2) a variation of high-intensity interval training (HIIT) we refer to as high intensity tactical training (HITT), to assess the dynamics of DNA methylation (DNAm) in blood and muscle derived from males (N=23) and females (N=31), over a 12-week period of training followed by a 4-week period of detraining, sampled at pre-exercise and acute time points, totaling 528 samples. Due to its rapid responsiveness to stimuli and its stability, DNAm has been known to facilitate regulatory cascades that significantly affect various physiological processes and pathways. We find that several thousand differentially methylated regions (DMRs) associated with acute exercise in blood, many of which are shared across males and females. This trend is reversed when comparing the baseline (pre-exercise) time points or post-exercise timepoints at the untrained state with those at the post-conditioned state. Here, muscle shows majority of DNAm changes, with most of those being unique. We also find several hundred “memory” DMRs in muscle that maintain the gain or loss of methylation after four weeks of inactivity. Comparing phenotypic measurements, we find specific DMRs that correlate significantly with mitochondrial function and myofiber switching. Using machine learning, we select a subset of DMRs that are most characteristic of training modalities, sex and timepoint. Most of the DMRs are enriched in pathways associated with immune function, cell differentiation, and exercise adaptation. These findings reveal mechanisms by which exercise- and training-induced epigenetic changes alter immune surveillance, mitochondrial function, and inflammatory response, and underscore the relevance of epigenetic plasticity to health monitoring and wellness.

## Introduction

Following decades of research, the first official recommendation in 1995 required a minimum of 30 minute moderate-intensity physical activity (PA; equivalent to 3.0-5.0 metabolic equivalents (METs) per hour) on most days of the week to increase the health benefits ^21^. More detailed and in-depth studies have ensued since then and have led to periodic updates to these guidelines ^22^ that include health benefits specific to age-groups, intensity of activity, and timing, among others. Based on a recent survey of approximately 2 million people, it was found that 27.5% of the world’s adult population do not meet recommended PA standards^23^. Sedentary lifestyle (MET <= 1.5) is a leading risk factor for noncommunicable diseases, with 20-30% increased risk of death compared to people who are sufficiently active ^24^. Analysis of pooled data from six prospective cohort studies involving 654,827 individuals have provided quantitative evidence of increased life expectancy across a range of activity levels and body mass index (BMI) groups, including an increase of 4.5 years following the minimal World Health Organization (WHO) recommended PA (300-449 minutes per week) and normal BMI (18.5-24.9) ^25^. Strong evidence has also been found for dose-response relationship to cardiorespiratory health, metabolic health and cancer risk ^26,26^.

Various exercise prescriptions offer distinct health benefits. For example, Moderate-Intensity Continuous Training (MICT) increases aerobic capacity, metabolic health, and endurance ^11^, moderate intensity, continuous endurance exercise program followed by a standard resistance training protocol (TRAD) ^10^ increases muscle strength, and metabolic rate, low-load resistance training with blood flow restriction (BFR) promotes muscular adaptations ^20^, or High-Intensity Interval Training (HIIT) increases VO₂max, mitochondrial function, and metabolic regulation ^13^. Integrating explosive bodyweight movements and exercises with external resistance, executed with minimal rest engages more muscle groups and optimizes balance and coordination, aimed to provide combined endurance and strength/power benefits. This prescription is widely used among athletes and in the military, including the US Marine Corps’ High-Intensity Tactical Training (HITT)^27^. Such tangible phenotypic, physical and wellness outcomes associated with exercise has been found to be a result of alterations that occur at the molecular level, within cells, that in turn affect gene expression profiles, release of cytokines, ATP turnover, and regulation of DNA damage among others ^28–32^.

One such molecular alteration is the covalent addition of a methyl group to the cytosine residue of DNA converting it to 5-methyl cytosine (5mC) ^33^. DNA methylation (DNAm) is a dynamic, yet stable epigenetic modification. Switching of the methylation status of cytosine residues at single sites - differentially methylated sites (DMS) - or as a group - differentially methylated regions (DMR) - has been known to cause dramatic changes in gene expression. It is a natural phenomenon in the context of cellular development, cell identity maintenance, differentiation, and aging. Hypermethylation of promoter CpG islands is generally associated with transcriptional silencing of tissue-specific genes while hypomethylation in the gene body is linked to gene activation ^34–37^. External stimuli, such as chemical and biological exposures ^38^, have also been known to change the methylation status of the cytosines in the genome. DNAm and histone modifications are found to be the most significant epigenetic changes linked to skeletal muscle transcriptional responses to exercise that mediate exercise adaptations ^39^. The association of DNAm and epigenetic memory of exercise-induced skeletal muscle hypertrophy was found in chronic resistance exercise ^31^. It has also been shown that the exercise responsive methylation changes are also associated with a reduced risk of chronic disease ^40^. To determine the findings of molecular changes resulting from physical activity and transform this knowledge into health outcomes, it is imperative to consider sex as a biological variable. In recent years, studies including both males and females have been designed with this necessity in mind ^41,42^.

As part of the Precision High Intensity through Epigenetics (PHITE) study ^27^, we assessed the dynamics of DNAm in response to acute exercise bout (acute effect), adaptation to chronic exercise (training effect), and inactivity (detraining effects) in the context of two training modalities (HITT and TRAD) and biological sex (male and female). Both exercise prescriptions lasted 12 weeks with training performed 3 days per week followed by 4 weeks of detraining, for a total study length of 16 weeks. Phenotypic measurements are recorded at relevant time points during this course. Sample preparations from venous whole blood and skeletal muscle biopsies (vastus lateralis) for this study were collected at five time points – (1) resting baseline before any training (w0pre) and (2) at three hours post-acute exercise (w0h3), (3) resting baseline after 12 weeks of training (w12pre) and (4) at three hours post-acute exercise (w12h3), and (5) after four weeks of detraining (w16rst). The overall study design^27^ is shown in figure 1A-C, metadata in Supplementary Table S1 and assay summary in Supplementary Table S2. A subset of these samples, specifically at the week 0 acute exercise were used for analyze the methylome using bead array, transcriptome, and metabolome by our collaborators ^41^.

**Figure 1:**
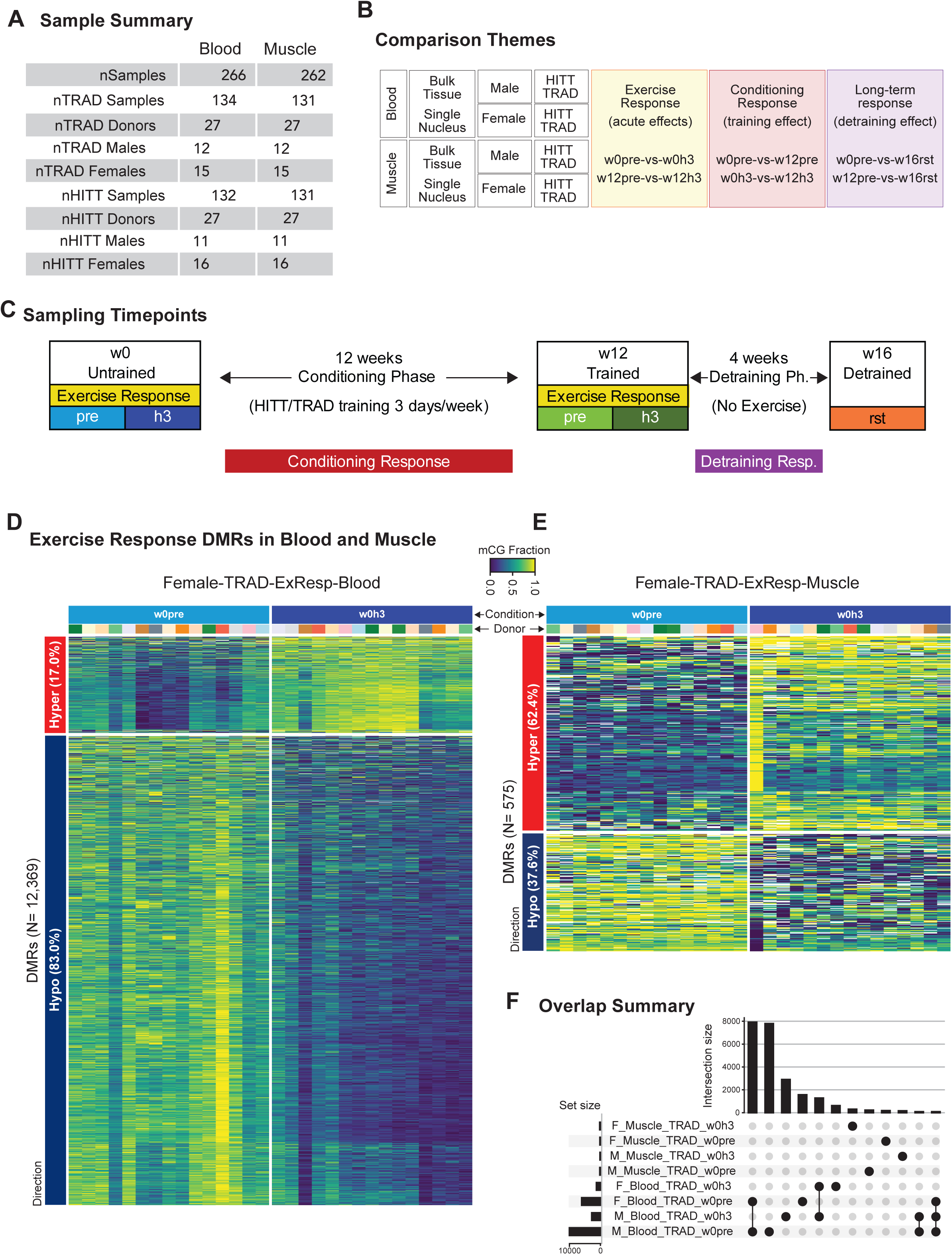
Study design and Exercise Response. (A) Overview of blood and muscle samples used in the study separated based on exercise dosage and sexes. The participants were stratified based on sex and an equal number was assigned to either the TRAD or HITT group. (B) Overview of major comparisons being tested. Within each tissue, we group the donors by sex and exercise dose to perform three themes of comparisons - (1) Exercise Response, where the acute effects are explored by comparing the methylation signals before exercise and three hours post exercise; (2) Conditioning Response, where the effect of training over the 12-week period is tested by comparing the baseline (pre-exercise) timepoint at week 0 (pre-training) and at week 12 (post-adaptation); (3) Long-term response, where we explore if any of the methylation changes persist after a four-week period of no exercise. Here we compare the baseline time points from week 0 and week 12 with the week 16 sample (sampled at a pre-exercise timepoint). (C) The sample collection timepoints are shown along with the training state and the comparison themes. (D) The heatmap is an example depicting the gain and loss of methylation at specific cytosine sites across the genome (rows) and how they are consistent across replicates (columns) of the comparison groups (w0pre and w0h3) in the Female-TRAD group in blood. Here, majority (83%) of the DMRs lose methylation at the acute timepoint. On the contrary, in muscle (E), most of the DMRs (62%) gain methylation at the acute timepoint, whereas the number of DMRs are considerably lower. (F) The Upset plot shows the number of hypo-methylated samples in each of the exercise comparison groups. Several thousand DMRs in blood are shared across the comparisons, whereas the DMRs in muscle are unique (shared DMRs less than 200 are not shown).

## Results

We explored the association between physical activity on DNAm changes using three major themes - (1) acute exercise effects, defined as the changes occurring at three hours (h3) post-exercise at both w0 and w12; (2) training adaptation, defined as changes due to the 12 weeks of training (across w0 and w12, either at pre-exercise or at three hours post exercise), and (3) The detraining effect, defined as changes that are seen at resting baseline levels after a four week period of no training or rest (rst) compared to the methylation patterns at untrained or trained state (i.e., w0pre vs w16rst or w12pre vs w16rst).

### DNA Methylation Dynamics of Acute Exercise

Overall, we observe 25 times the number of differentially methylated regions (DMRs) in whole blood (average 14,666 regions, std. dev. ± 4,296) compared to muscle (average 597 regions, std. dev. ± 65) as a result of acute exercise (p < 0.001; Welch’s t-test; Fig. 1D-E). We further explore the functional relevance of these DMRs, especially with the knowledge that inflammatory responses, T-cell migration, apoptotic pathways, and hypoxic conditions result in changes more observable in blood at the acute timepoint.

In blood, males had a greater number of acute response DMRs than females (average of 16,547 and 12,785, respectively). Chronic training reduced the number of acute response DMRs from an average of 17,835 at w0h3 to 11,447 at w12h3, suggesting training adaptations. The exercise modalities (HITT and TRAD) did not show a striking difference in the number of DMRs.

In muscle, females have slightly more number of DMRs than males (N=622 and 572 for females and males respectively). The number of DMRs in the HITT group were higher (N=639) compared to the TRAD group (N=555) (Supp. table S3).

The DMRs in blood associated with acute exercise are mostly shared across sex and exercise prescription (N=8,083 at w0 and N=4,486 at w12), whereas the DMRs in muscle are mostly unique. At w0 phase in blood, the DMRs shared for females across prescriptions are the least (N=286), compared to the shared DMRs across prescriptions for males (N=2,593), which highlights the sex-specific response to the intensity of acute exercise (Fig 1D, 1E). Females that performed HITT had a DMR signature that overlapped with males (both HITT and TRAD; N=3,115) much more than the females that performed TRAD (N=678). More DMRs are shared across males and females in the HITT group (N=723) compared to TRAD (N=462). At w12, the number of shared DMRs across the four groups was reduced from N=8,083 to 4,486, even though the number of DMRs in groups did not change as much. The same trends were observed for the shared DMRs between the groups as observed at w0. Comparing all acute phase DMRs across w0 and w12, we find 4,078 DMRs shared across all groups (Fig 1F). Male-TRAD and Male-HITT groups have the most number of unique set of DMRs (N=4,343 and 3,486, respectively). Male-TRAD and -HITT groups also share the greatest number of DMRs (N=1,428). Fig. S1 summarizes the breakdown of DMRs in response to acute exercise, with Fig. 1F showing TRAD as an example.

DMRs can be classified as regions that either gain (hyper) or lose (hypo) methylation. Within the 17,126 DMRs in blood for Female-HITT, ∼80% of them (N=13,747) are hypomethylated at the acute time point in the untrained state (w0h3). This proportion is similar at the trained time point as well (10,732 hypomethylated DMRs at w12h3 out of 13,146 DMRs). We also see the same trend for all other categories of Female-TRAD, Male-HITT, Male-TRAD, at both w0 and w12.

In muscle, the number of DMRs were much smaller and so were the proportions of hypomethylated DMRs at the three hours post-exercise timepoint. For the Female-HITT category, only around 60% of total DMRs are hypomethylated following exercise at w0 (N=386/641) and only 41% of DMRs are hypomethylated after acute exercise at w12 (N = 272/652). For the Female-TRAD group, 37% (217/575) and ∼47% (287/614) of DMRs are hypomethylated after acute exercise at w0 and w12, respectively. For the Male-HITT group, ∼50% DMRs are hypomethylated after acute exercise at w0 and w12 (N=308/614 and 335/643, respectively). For the Male-TRAD group, ∼60% DMRs are hypomethylated after exercise at w0 (N=271/454) with only ∼36% hypomethylated at w12 (N=205/572).

### DNA Methylation Dynamics of Training Adaptation

One of the major objectives of this work was to study epigenetic dynamics of training adaptation to HITT or TRAD prescriptions (Fig 2). Consistent supervised training is expected to create stable patterns of physiological and molecular changes. It has been shown that endurance training leads to molecular adaptations in both skeletal muscle and blood, such as mitochondrial function, capillary density and metabolic efficiency^44^.

**Figure 2:**
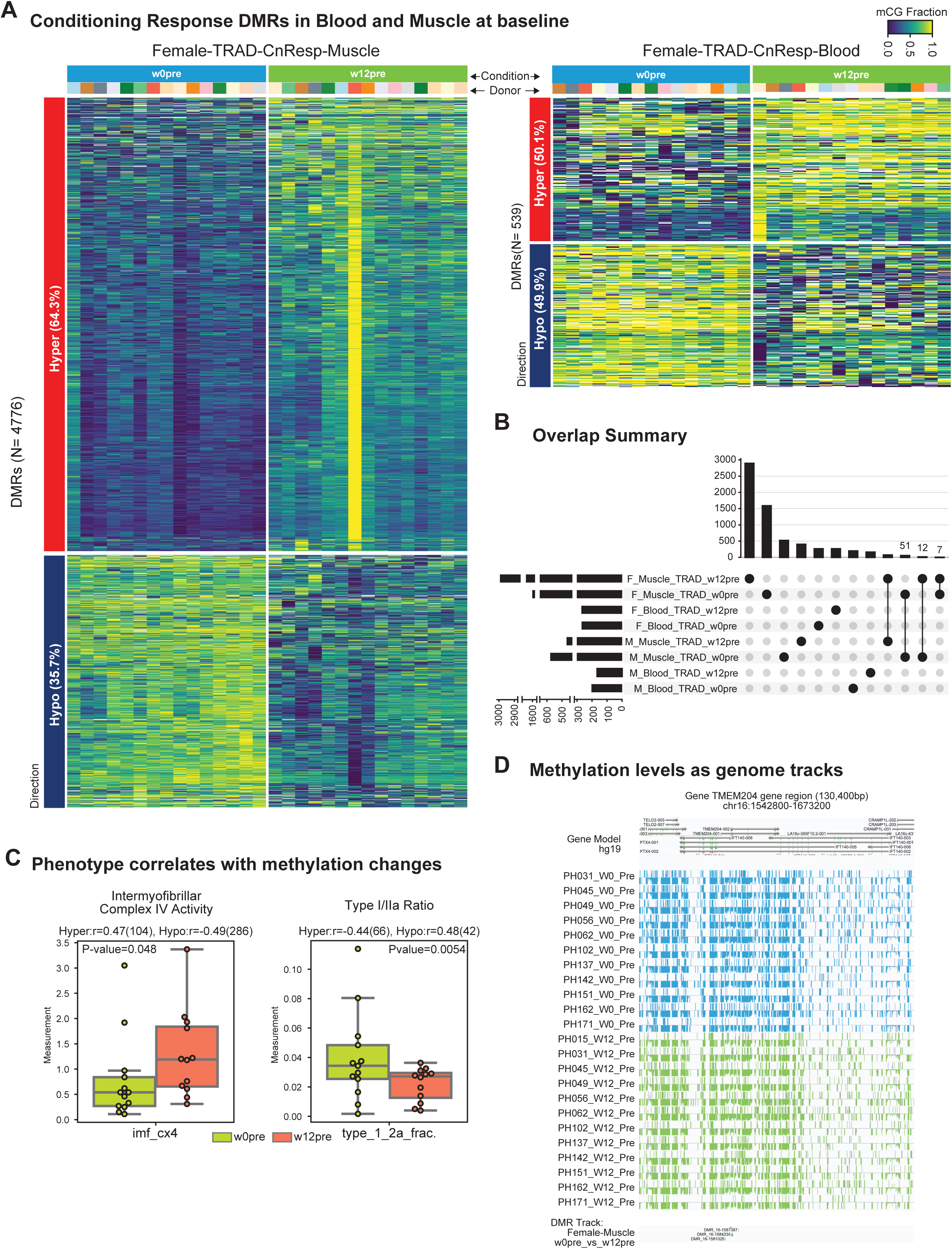
Conditioning Response. (A) The heatmap is an example comparing the methylation levels at baseline (pre-exercise) timepoint before and after the 12-week training in the Female-TRAD group. In contrast to the DMRs found in blood at acute phase, ∼64% of the DMRs gain methylation after the consistent training period. Although there are a few outliers, the replicate consistency (columns) and direction of change is maintained across the genomic sites (rows). In contrast, much fewer DMRs are identified in blood at the baseline timepoint. (B) The Upset plot shows hypo-methylated samples in each of the conditioning response comparisons. Very few DMRs are shared in this comparison theme. (C) We identify several conditioning response DMRs that correlate well with the phenotypic measurement. The examples shown are for intermyofibrillar complex IV activity which is elevated at week-12 and is correlated with 104 hyper-methylated sites and 286 hypo-methylated sites. Also shown as an example is the ratio of TypeI/Type IIA myofibers which reduces at w12. This is correlated to the methylation level from 66 hypermethylated sites and 42 hypomethylated sites. (D) An example of a differentially methylated region is shown as a browser track, focusing on the TMEM204 gene (Fig 2D).

In skeletal muscle, the number of DMRs that changed as a result of training were 1,999 for Female-HITT (∼50% hypermethylated), 4,776 for Female-TRAD (64% hypermethylated), 1,353 for Male-HITT (65% hypermethylated), and 1,074 for Male- TRAD (45% hypermethylated), These are higher than those observed in blood (N=323 for Female-HITT, 539 for Female-TRAD, 541 for Male-HITT, and 369 for Male-TRAD). It is observed that differential methylation is greater in females compared to males, in particular in the Female-TRAD group (4 times greater compared to Male-TRAD) (Fig 2A).

After acute exercise, chronically trained muscle shows 1,172 DMRs in Female-HITT (60% hypermethylated), 1,370 DMRs for Female-TRAD (33% hypermethylated), 1,957 for Male-HITT (57% hypermethylated), and 3,295 for Male-TRAD (66% hypermethylated). The number of DMRs in blood are much lower (N=376 for Female-HITT, 437 for Female- TRAD, 397 for Male-HITT, and 543 for Male-TRAD). Here, we see the Male-TRAD group having ∼2.5 times more DMRs compared to the Female-TRAD group. An example for this is the gene TMEM204 (Transmembrane Protein 204) was found to be a differentially methylated gene in the Female-TRAD group comparing w0pre_vs_w12pre as well as harboring several DMRs (Fig 2D). This gene was also found to be associated with exercise training in adipose tissue which also studied DNA methylation levels and gene expression ^45^. TMEM204 is involved in the regulation of VEGF receptor signaling pathway (which is also an enriched function in conditioning response). It is also known to be involved in smooth muscle cell differentiation.

We overlap DMRs that are within 1kb of each other on a sliding window and find 70 DMRs shared across the muscle samples (Fig 2B; Supp. Fig. S1).

### DNA Methylation Dynamics of Detraining

We then explored the changes in DNAm associated with 4 weeks of inactivity that occurred after 12 weeks of supervised training (w16rst). In blood, compared to pre-training baseline (w0pre), we see 406 DMRs (36% hypermethylated) in the Female-HITT group, 467 for Female-TRAD (48% hypermethylated), 500 DMRs Male-HITT (37% hypermethylated), and 448 for Male-TRAD (39% hypermethylated). We observed much greater numbers in skeletal muscle. N=1,976 for Female-HITT (67% hypermethylated), 2,053 for Female-TRAD (66% hypermethylated), 6,553 for Male-HITT (68% hypermethylated), and 969 for Male-TRAD (51% hypermethylated). The Male-HITT group has the largest number of DMRs (∼60% more than other groups). Out of the 68% of DMRs that are hypermethylated in the Male-HITT group compared to untrained baseline(N=4,431), only 696 of these are shared with any other group.

While comparing w16rst to the w12pre group, we see similar trends in blood (N=433 for Female-HITT, 454 for Female-TRAD, for 438 Male-HITT, and 402 for Male-TRAD) and muscle (N=591 for Female-HITT, 660 for Female-TRAD, 1,174 for Male-HITT, and 452 for Male-TRAD) (Supp. Fig. S2).

### Differentially Methylated Genes

Differentially methylated gene (DMG) analysis revealed several genes consistently altered across various experimental conditions, reflecting significant changes in gene-body DNAm associated with acute exercise, training adaptation and detraining. Out of 48 total comparisons, eight comparisons had at least one DMG with an adjusted p-value less than 0.05. Of these eight comparisons, five DMGs were exercise responsive in blood, two were training responsive in muscle, and one was a de-training response in muscle. Of these DMGs, we noted several associated with the immune system, metabolism, muscle adaptation, and cancer, using the Database of Archived Variations of Intracellular Dynamics (DAVID)^46^.

ANKRD18A was a DMG in four out of five blood comparisons. Interestingly, this gene displayed decreased methylation post-exercise, suggesting exercise-induced demethylation. Prior research has demonstrated that hypermethylation of ANKRD18A serves as a biomarker for lung cancer and could potentially mediate disease pathogenesis ^46^. The observed exercise-associated hypomethylation of ANKRD18A may imply protective epigenetic adaptations.

Another biologically relevant DMG identified was G0S2, known for its critical role in lipid metabolism and regulation of adipocyte function. Differential methylation of G0S2 following exercise could influence lipid storage and mobilization, potentially contributing to beneficial metabolic adaptations driven by physical activity.

### Phenotype Correlations with DMRs

Our studies complement the comprehensive phenotyping measurements conducted under the parent clinical trial^27,41^. The main objective was to assess the changes due to chronic training. 14 categories (such as Wingate measurements, DXA scan, muscle histology, among others.), conducted at w0, w12, and w16rst. We identified 2,598 hyper- and 2,655 hypo-DMRs whose methylation levels were significantly correlated with phenotypes showing differential measurements (Wilcoxon test p-value ≤ 0.05). For example, the intermyofibrillar complex IV activity (’imf_cx4’) measurement significantly increased (p-value = 0.048) in the w12pre group compared to the w0pre group and the ratio of Type I to Type IIA myofiber phenotype showed a significant difference between the w0-pre and w12-pre groups (p-value = 0.0054, Figure 2C). In this specific comparison (w12pre vs. w0pre), we found 104 hyper- and 286 hypo-DMRs. The median Spearman correlation coefficient between the ’imf_cx4’ measurement and the methylation fraction of these hyper- and hypo-DMRs were 0.47 and -0.49, respectively. This indicates that increased methylation in hyper-DMRs is associated with higher ’imf_cx4’ measurements in the w12pre group compared to the w0pre group, while decreased methylation in hypo-DMRs shows the opposite trend. Comparing w12pre versus w0pre, we identified 66 hypermethylated and 42 hypomethylated DMRs, with median Spearman correlation coefficients of -0.44 and 0.48, respectively. Summary of correlation of each marker is provided in Supplementary Table S4.

### Longitudinal Variation in DNA Methylation

For time course analysis, DNAm data were normalized to each participant’s initial methylation, which was treated as a personalized “basal” methylation level. To identify methylation sites that change significantly over time, a B-spline was fitted to each DMS. When analyzing DNAm at rest over three time points (w0 pre, w12pre, w16rst), a total of 61,134 Longitudinally Variable Methylation Sites (LVMS) were identified as significantly changing over time across sex and training regimen. Notably, females had far more LVMS than males in muscle tissue (Supp. Fig S4i.). While females who had complete the TRAD prescription saw more LVMS than their HITT counterparts, the reverse was observed for males. In male muscle, those that had undergone the HITT exercise saw more LVMS than their TRAD counterparts.

To further characterize the 61,134 sites, we clustered them in two stages. The first stage used KMeans to cluster on each spline’s median values. The silhouette score was used to determine the optimal k parameter (Supp. Fig. S4 ii., k = 26). Spline clusters were then characterized by their sex and training group as well as their CG/CH composition (Supp. Fig S4 iii.). These clusters were also grouped together in higher order clusters by each cluster’s largest time course change, whether it was between w0 and 12 (early) or w12 and w16 (late), as well as if the change resulted in a positive or negative change in methylation (up/down). The sole Male HITT dominated clusters were characterized by a stronger decrease in methylation sites during the detraining phase. While this higher order clustering approach may be highly interpretable, the resulting four higher order clusters do not always best capture the splines, except for the late decrease methylation category (Supp. Fig S4 iv. A). This was evident when performing a hierarchical clustering of each spline cluster’s median slopes. (Supp. Fig. S4 iv. b). We then used hierarchical ordering to determine seven higher order clusters (Fig. 3A, Supp Fig S4 v. A, B).

**Figure 3:**
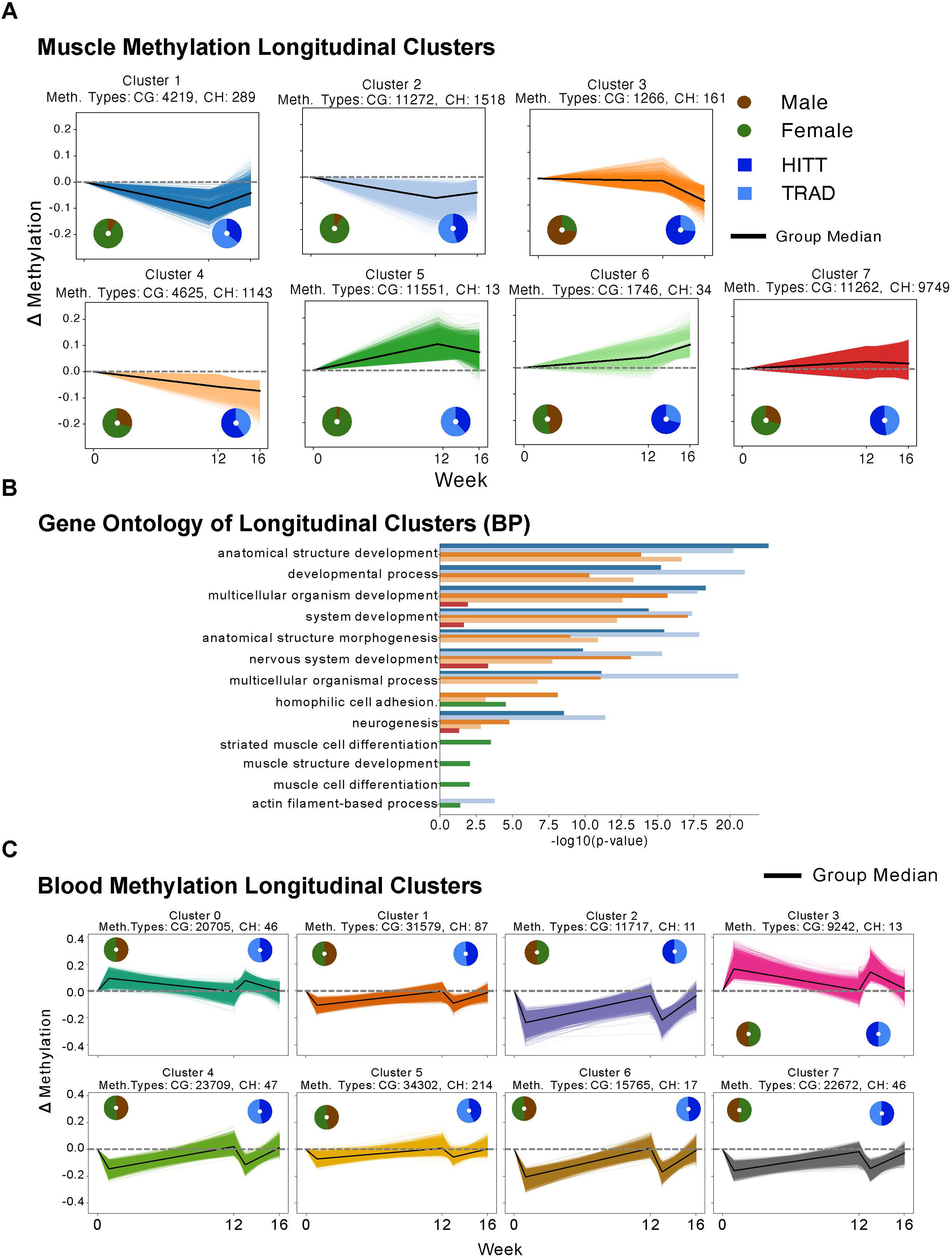
Longitudinally Variable Methylation Sites. (A) Line plots of longitudinal clusters in muscle tissue across pre-exercise and rest phase. Accompanied with pie charts of sex and exercise composition per cluster. B) Bar plots of top 5 gene ontology (BP) terms per cluster. C) Line plots of clustered splines with five time points in blood.

Methylation sites within clusters were then annotated with gene regions and gene ontology was performed across three GO domains of Biological Process (BP), Cellular Component, (CC), and Molecular Function (MF) with a background gene set of all genes across clusters (Fig 3B, Supp. Fig S4 vi. A,B). Clusters 1-4 appeared to be strongly interrelated in their biological functions. These three clusters were all significantly enriched for developmental processes, transcriptional regulation, and intercellular communication. These methylation sites likely correspond to not just the genes involved in intercellular growth processes, but also their transcriptional regulators as well. Clusters 5 and 7 differ significantly in their processes compared to clusters 1-4. The sites in clusters 5 and 7 correspond to cytoplasmically localized genes that are directly involved in muscle growth as well as protein and ion binding. In other words, clusters 1-4 likely correspond to intercellular regulation for muscle and nerve development, whereas clusters 5 and 7 correspond to intracellular protein processes for muscle growth (Fig 3B). We then sought to determine if the methylation sites between clusters were also correlated positionally, not just functionally. We binned the number of LVMS in 500 bp width bins throughout the genome and normalized the counts to the size of each cluster (Supp. Fig S4 vii. A, B). We then performed Principal Component Analysis (PCA) on these normalized, binned, counts and correlated the clusters’ normalized counts against each other. (Supp. Fig S4 viii. a, b). These positional correlations largely agreed with our function relationships. Clusters 1-4 had less variance amongst each compared to each other than clusters 5 and 7. Clusters 5 and 7 were correlated with each other, though they did show significant separation along the first two principal components. This difference might be because cluster 7 has far more CH methylation than cluster 5. Clusters 1-4 were also mostly correlated with each other, with the notable exception of cluster 3, the only male dominated higher order cluster. We then analyzed which sites were the most specific for clusters using a metric that incorporated both entropy for each site and the difference between absolute normalized counts (Supp. Fig S4 ix A, B). Our results show that clusters 3 and 6 have more highly cluster-specific sites compared to other clusters. We annotated the methylation sites with genes as well and found several genes of note that were highly specific to cluster 3, namely GDNF, RNF219-AS1, and LINC00982. Notably, GDNF is strongly present in nervous system functions while RNF219-AS1, and LINC00982 are non-coding RNAs.

We then applied our spline modeling analysis on all five time points focusing on the blood samples. Consistently, the largest change in blood was the change from pre- to post-acute exercise (Fig 3C). This exercise response induced changes in both hyper and hypomethylation, and these changes remained consistent after chronic training, indicating that within our blood samples, exercise would consistently alter methylation regardless of the timepoint of the study.

### Differential Transcription Factor Motif Usage

Distinct families of transcription factor (TF) motifs are enriched in DMRs of different conditions, even within TF families. For example, the SOX7 motif was enriched in DMRs losing methylation in Male-HITT group whereas the SOX2 motif was enriched in DMRs that lose methylation in Male-TRAD group (w0h3 vs w12h3) in muscle.

In muscle, the TF motif enrichment is mostly at sites losing methylation and is shared with both the chronic training response and following detraining, suggesting a sustained regulatory memory. We mainly see three groups of TFs enriched (Fig 4A).

**Figure 4:**
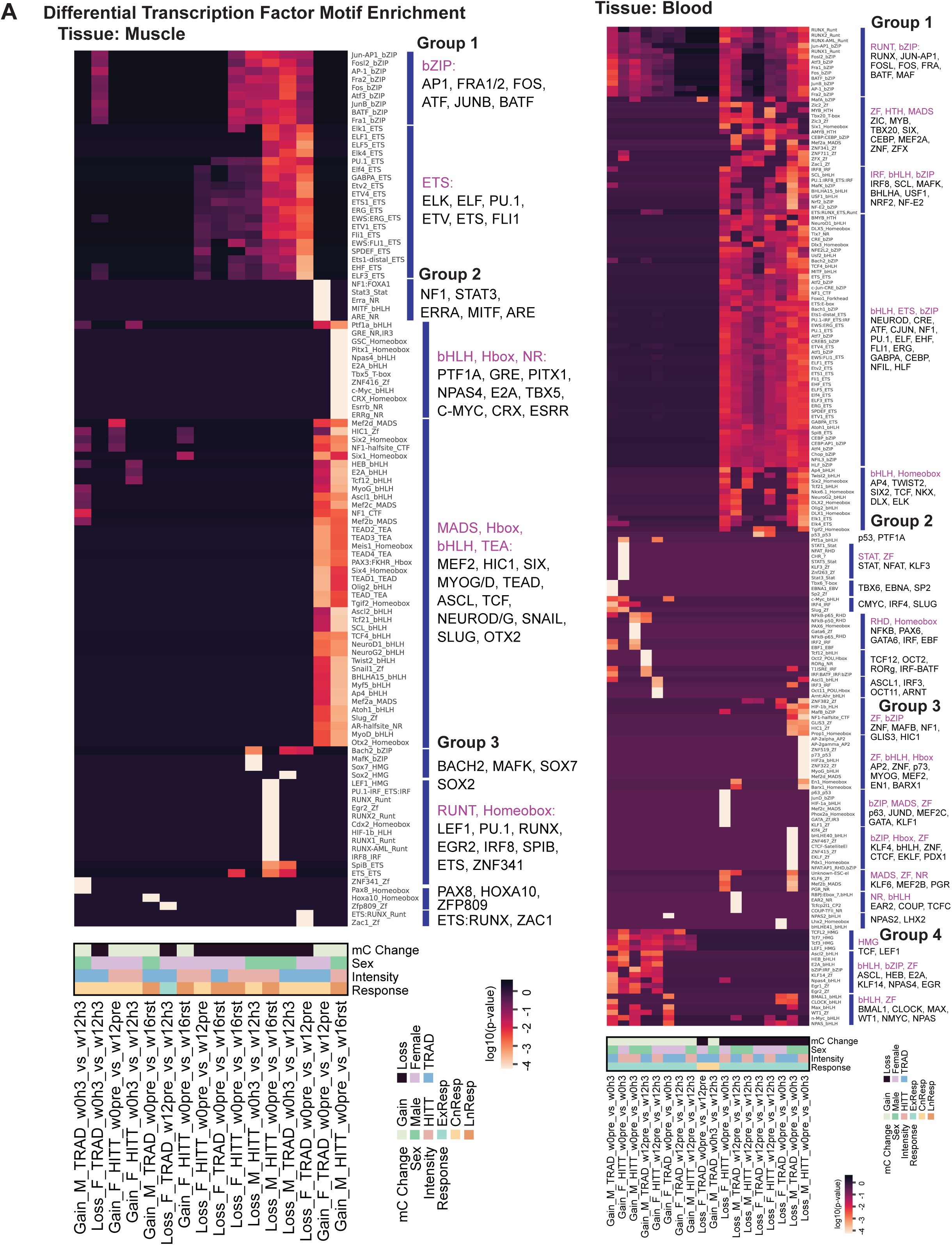
Motif enrichment at DMRs. (A) Three major groups of transcription factor motif enrichment can be seen which align to DMRs that lose methylation after the training period (measure at the acute timepoint), DMRs that gain methylation after the training (measured at the baseline), and a set (Group 3) of TFs (RUNT family) enriched in sites that maintain the loss of DNA methylation even after detraining. Similarly, for blood (B) we see four groups of TFs enriched in DMRs that gain methylation at acute time point or lose methylation. These TFs are known to affect distinct pathways.

Group 1 TFs can further be sub-grouped based on various TF families. Overall, they are enriched in DMRs that lose methylation in the following comparisons: Female-HITT chronic adaptation at pre-exercise timepoint, Female-TRAD w0pre_vs_w16rst, Female- HITT w0pre_vs_w16rst, Male-HITT w0h3_vs_w12h3, Male-HITT w0pre_vs_w16rst, Male-TRAD w0h3_vs_w12h3, and Female-TRAD w0pre_vs_w12pre. These TFs can further be grouped based on the TF families. The bZIPs, AP1 (FOS, JUN, FRA, BATF) are master regulators of stress and mechanical signals in muscle ^48^. The ETS family of TFs (ELK, ELF, PU.1, ETV, ETS, FLI1) are involved in cell proliferation and differentiation^48^.

Group 2 TFs are enriched in DMRs that Gain methylation at w12pre and w16rst timepoints in the Female-TRAD w0pre_vs_w12pre and Male-HITT w0pre_vs_w16rst comparison respectively. These TFs can also be distinctly classified as bHLH, H-box, NR TFs (NF1, STAT3, ERRA, MITF, ARE, PTF1A, GRE, PITX1, NPAS4, E2A, TBX5, C-MYC, CRX, ESRR). They play vital roles in muscle adaptation and recovery ^50^, mitochondrial biogenesis ^51^, and differentiation. The MADS, bHLH, TEA family (MEF2, HIC1, SIX, MYOG/D, TEAD, ASCL, TCF, NEUROD/G, SNAIL, SLUG, OTX2) of TFs regulate metabolism ^52^ and satellite cell differentiation.

Group 3 TFs are very specific to certain DMRs only. BACH2, MAFK and SOX7 TF motifs are enriched only in DMRs that lose methylation in w12h3 in the M_HITT_w0h3_vs_w12h3 comparison. These TFs are involved in angiogenesis and antioxidant functions; while SOX2 is enriched only in the TRAD group of the same comparison. SOX2 is a key regulator of stem cell (satellite cell) pluripotency. The RUNT, homeobox family TFs (LEF1, PU.1, RUNX, EGR2, IRF8, SPIB, ETS, ZNF341) are involved in muscle regeneration ^53^, neuromuscular junction remodeling ^54^, and immune response regulation ^55^.

PAX8, HOXA10, ZFP809 TF motifs are enriched specifically in DMRs that gain methylation at w12h3 in the Male-TRAD w0h3_vs_w12h3 comparison.

In blood, the motif enrichment is grouped by DMRs that gain or lose methylation at acute phase (exercise response) suggesting a regulatory cascade that responds to stimuli. We see four major groups of TF motif enrichment (Fig 4B).

In Group 1 we see TFs enriched in DMRs that lose methylation especially at the acute time point (h3) after chronic training. This group includes TFs such as RUNX1 involved in hematopoesis, AP1 complex (FOS, JUN, FOSL, FRA) regulating genes in response to stress and cytokine release, BATF which is crucial for the Th-cell differentiation and inflammation ^56^ ^57^, MAF family TFs are also involved in immune responses. MYB is involved in regulating cell proliferation. TBX20 is known to contribute to cardiovascular adaptations ^58^. The CEBP TF regulate immune responses and metabolism. MEF2 TF and its interaction with HDAC5 is known to increase metabolic capacity of the skeletal muscle, which is increased due to exercise ^59^. ZFX and the other zinc-finger protein TFs are essential for hematopoietic stem cell maintenance. BHLHA, USF1, NRF2, NF-E2 TFs are involved in oxidative stress responses and metabolic adaptations. NEUROD, CRE, and ATF regulate neuronal an metabolic gene expression which is influenced by exercise, contributing to neuroplasticity and metabolic heath ^60^. PU.1, ELF, EHF, FLI1, ERG, GABPA TFs are involved in vascular development. NFIL, HLF, AP4, TWIST2, SIX2, TCF, NKX, DLX, ELK also regulate various genes involved in development and immune responses.

In Group 2, p53 motif is specifically enriched in the DMRs that gain methylation in the Female-HITT w0pre_vs_w0h3 group. P53 is known to play a crucial role in regulating cell cycle and apoptosis. Exercise-induced stress can activate p53 pathways ^61, 62^ ^63^. STAT, NFAT and KLF3 are specifically enriched in DMRs that gain DMRs in the Male-TRAD w0pre-vs-w0h3 comparison. STAT is involved in cytokine signaling ^64^, NFAT plays a crucial role in T-cell development ^65^, and KLF3 is implicated in the suppression of inflammatory responses ^66^. CMYC and SLUG is known to increase in response to exercise and influence stem cell activity ^67^. IRF4 is known to regulate immune cell differentiation ^68^. NFKB, PAX6, GATA6, IRF, EBF TF motifs are specifically enriched in DMRs that gain methylation at w0h3 in the Male-HITT w0pre_vs_w0h3 comparison. NFKB is a major TF that regulates immune responses, leading to production of cytokines and modulation of inflammation in the context of exercise ^69^. GATA6 regulates genes associated with vascular smooth muscle function ^70^. IRF is also a critical F involved in interferon production and immune response ^71^. EBF family TFs regulate B-cell development ^72^. TCF12, OCT2, RORg, IRF-BATF are specifically enriched in DMRs that gain methylation at w12h3 in Male-TRAD w12 acute exercise comparison. TCF12 mediates the development of both T and B lymphocytes ^73^. OCT2 is a TF that is involved in the humoral immune response ^74^. RORG is involved in Th17 cell differentiation ^75^. Interaction between IRF-BATF is known to regulate T-cell differentiation ^75^. The TFs ASCL1, IRF3, OCT11, ARNT are enriched in DMRs that gain methylation at w12h3 in Female-HITT w12pre-vs-w12h3 comparison. Exercise is known to regulate IRF3 production which leads to enhanced antiviral immunity ^76^.

In Group 3, we see an enrichment of zinc-finger TFs, MAFB, NF1, GLIS3, AP2, p73, MYOG, and MEF2, enriched in the DMRs that lose methylation at w0h3 in the Male-HITT w0pre-vs-w0h3 comparison. MFAB, which is expressed in myeloid cells, is a master regulator of moncytopoiesis. Exercise is known to modulate transcriptional regulator of monocytes ^77, 79^. AP2 is involved in vascular development. p73 is a member of the p53 family of TFs and is involved in apoptosis pathways ^80^. MYOG (Myogenin) is a myogenic regulatory TF involved in skeletal muscle development, especially in the differentiation of myoblast to mature muscle fibers ^81^. MEF2D is involved in regulating T follicular helper (T-fh) cells and is involved in IL-21 mediated humoral immunity ^81^. p63, JUND, MEF2C, GATA, KLF1 TFs are enriched specifically in DMRs that lose methylation at w0h3 in the Female-HITT w0pre-vs-w0h3 comparison. JUND modulates oxidative stress response and is implicated in apoptosis pathways ^83^. MEF2C is involved in muscle metabolism and adaptation to exercise ^84^. It also plays a key role in development of slow-twitch myofibers^85^. GATA is well known for its role in erythroid development ^86^. The TFs KLF4, bHLH, ZNF, CTCF, EKLF, PDX1, KLF6, MEF2B, PGR are specifically enriched in DMRs that lose methylation at w0h3 in Male-TRAD w0pre-vs-w0h3 comparison. KLF4 enhances mitochondrial function, contributing to improved metabolic adaptations ^86^. bHLH TFs are crucial during stress hematopoiesis induced by exercise ^88^. CTCF mediates interactions essential for hematopoiesis ^88^.

In Group 4, the TFs TCF, LEF1, ASCL, HEB, E2A, KLF14, NPAS4, EGR, BMAL1, CLOCK, MAX, WT1, NMYC, NPAS are enriched in DMRs that gain methylation at h3 in various comparison groups. These TFs are generally involved in T-cell development and immunity ^90^.

### Function Enrichment of DMR-associated Genes

For the acute exercise response, in blood (Fig 5A), specifically in the HITT group at w12, for genes associated with DMRs that gain methylation after exercise we observe enriched functions of T cell receptor signaling pathway, Adherens junction, Primary immunodeficiency, Cytokine-cytokine receptor interaction, Viral protein interaction with cytokine, HIF-1 signaling pathway, MicroRNAs in cancer, and Th1 and Th2 cell differentiation pathways.

**Figure 5:**
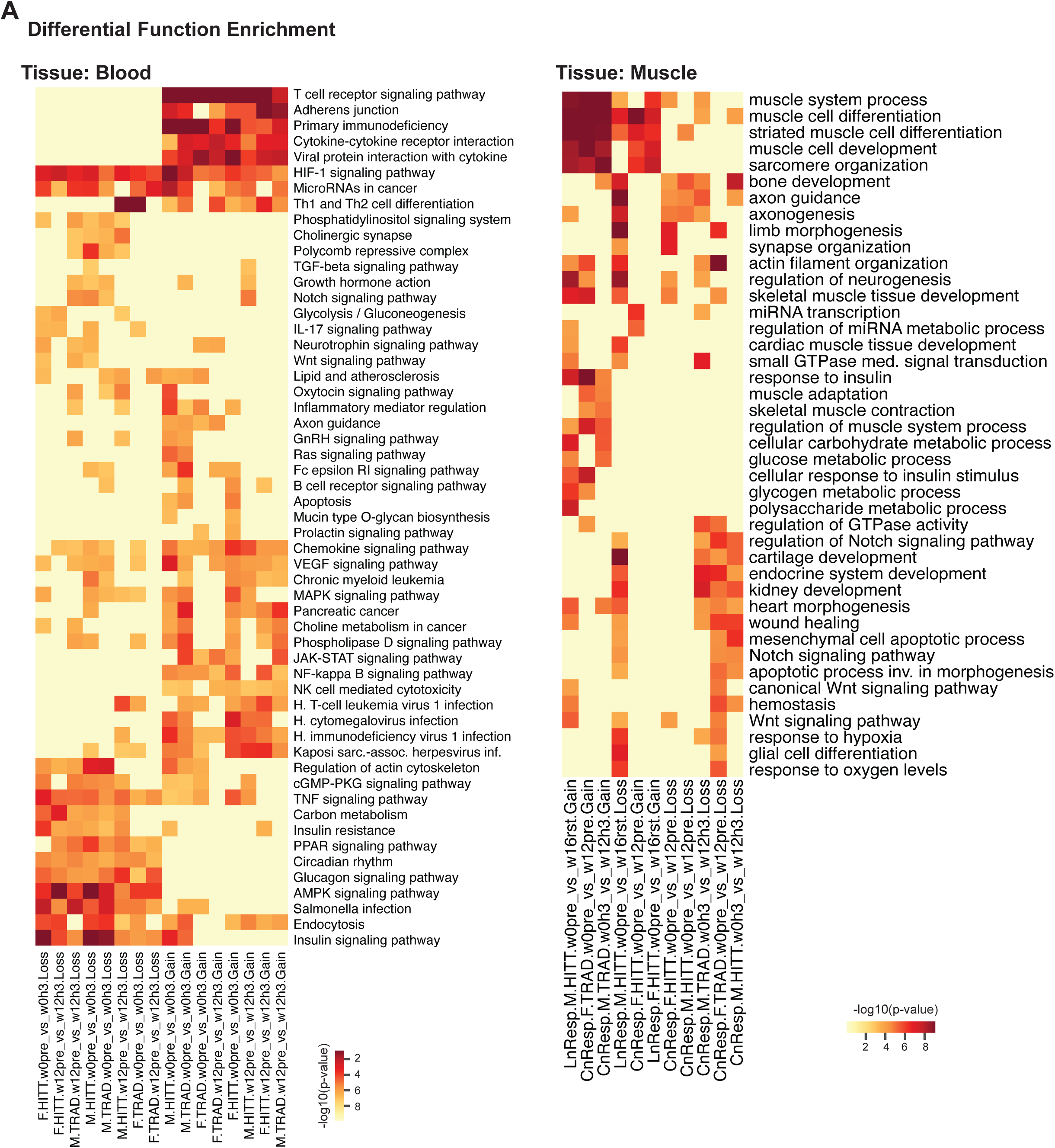
Function enrichment of genes associated with DMRs. Using a regulatory region defined around DMRs we identify genes associated with such sites and perform an enrichment analysis of biological processes. Based on semantic similarity, the major groups are identified in (A) blood and (B) muscle.

T-cell mobilization into the circulation during acute bouts of exercise, reduction of lymphocyte numbers and their redistribution to peripheral tissues as an immune surveillance mechanism has been well studied ^91^. Adherens junctions at cell junctions ensure the integrity of epithelial tissue and are key for muscle remodeling and adaptation to intense exercise ^92^. Primary immunodeficiency is also related to the above mentioned redistribution of immune cells and their migration. It is also known that acute exercise modulates myokines secretion from muscle ^93^. This effect is seen in the blood. The Viral protein interaction with cytokine receptor is also related to the above mentioned function ^94^. HIF-1 pathway simulates hypoxic conditions as a means of exercise adaptation ^95^.

Several microRNAs are associated with exercise response ^96^. The shift in Th1/Th2 (T-helper cells) balance has been observed in PBMCs at acute exercise, especially in trained individuals ^97–99^. This is clearly seen with the high enrichment of Th1 and Th2 cell differentiation pathway as most enriched in the w12 acute timepoint comparison. The other major pathways enriched in blood where we see gain in methylation following acute exercise are also related to immune activation and surveillance (Chemokine signaling pathway, Chronic myeloid leukemia, viral pathways), inflammation and repair (MAPK signaling and cancer pathways), and adaptation (VEGF signaling, Phospholipase D signaling pathway, NK cell mediated cytotoxicity).

For DMRs that lose methylation after acute exercise, we see enrichment of pathways involved in metabolic regulation (PPAR signaling, AMPK signaling and Insulin signaling pathways), immune function (TNF signaling, and infection), cellular adaptation (Regulation of actin cytoskeleton, Insulin resistance), and circadian rhythm.

For muscle, (Fig 5B), we see enriched pathways in DMRs from both conditioning response and long-term response. In both these responses, DMRs that gain methylation are enriched in muscle differentiation (muscle differentiation, sarcomere organization, actin filament organization, skeletal muscle tissue development), adaptation (skeletal muscle contraction, regulation of muscle system process, regulation of neurogenesis, response to insulin).

Pathways enriched for genes associated with DMRs that lose methylation mainly include repair pathways and satellite cell differentiation such as Regulation of GTPase activity, regulation of Notch signaling pathway, cartilage development, endocrine system development, kidney development, heart morphogenesis, wound healing, mesenchymal cell apoptotic process, Notch signaling pathway, apoptotic process inv. in morphogenesis, canonical Wnt signaling pathway.

### Minimal Set of Most Informative DMRs

This data offers the unique advantage of studying the effects of several parameters, such as exercise intensity, training prescription, and detraining in two tissues and two biological sexes. We have defined several hundred to thousands of genomic regions that respond to these combinations in terms of gain or loss of DNAm. However, identifying a minimal set of genomic regions that can serve as a specific signature responsive to training modality and sex provides a useful method to help understand the biological effects discussed earlier. Using these features, we developed a predictive model to distinguish combinations of training prescription and sex (Fig. 6A). The robustness of these DMR subsets enables the model to achieve an average accuracy of 0.88 across comparisons and tissues. The subset DMRs (features) are selected such that the donor effect is minimized. This allows for a more general and universal application of the models. Subsets of 0.1% to 43% DMRs can predict the comparison group with an AUROC greater than 0.95 (Fig. 6B, 6C, Supp. Table S6; Supp. Fig. S3).

**Figure 6:**
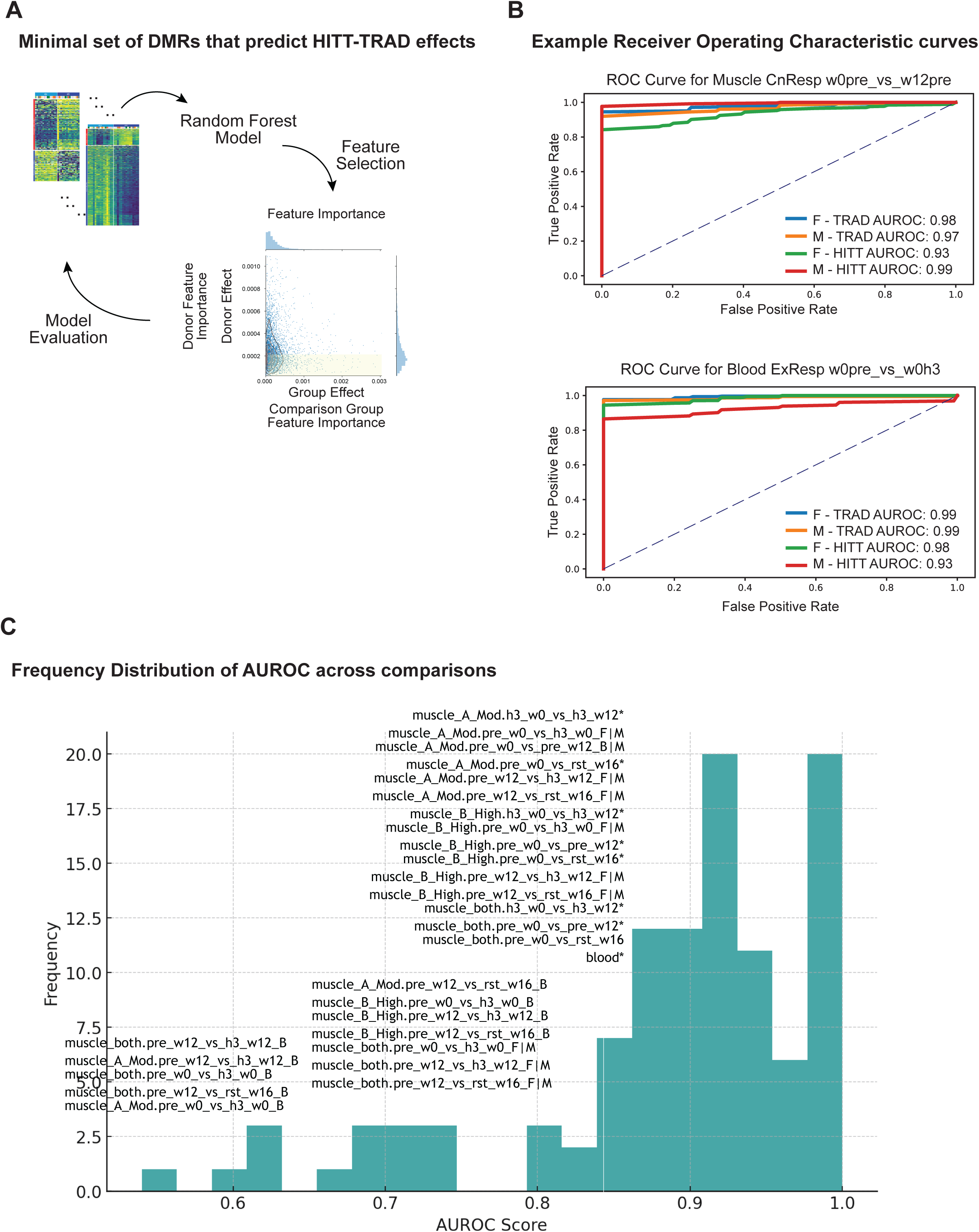
Machine Learning Approach to identify the most informative DMRs (A) Overview of the iterative implementation of the random forest model (B) example distribution of prediction power of the most informative DMRs shown for Muscle (top) in condition response and Blood (bottom) for exercise response for each of the groups. (C) the frequency distribution histogram of AUROC values for each of the 48 comparison groups are shown.

### Differential Methylation at Single-Nuclei Resolution

Single-nuclei analysis using PBMCs enabled us to account for the imbalance of cell composition before and after exercise in whole blood (Fig 7A). We see a marked increase of monocytes and Th-Memory cells at the acute time point, whereas Th-naive, Tc-Memory, NK-cell and B-cell counts drop at the acute time point. We see a considerable increase of NK-cells at the detraining timepoint. We further cluster the nuclei based on the global mCG fraction (Fig 7B, 7C) and further annotate them based on the antibody signals for each of the eight agranulocyte populations that were selected (Fig 7D).

**Figure 7:**
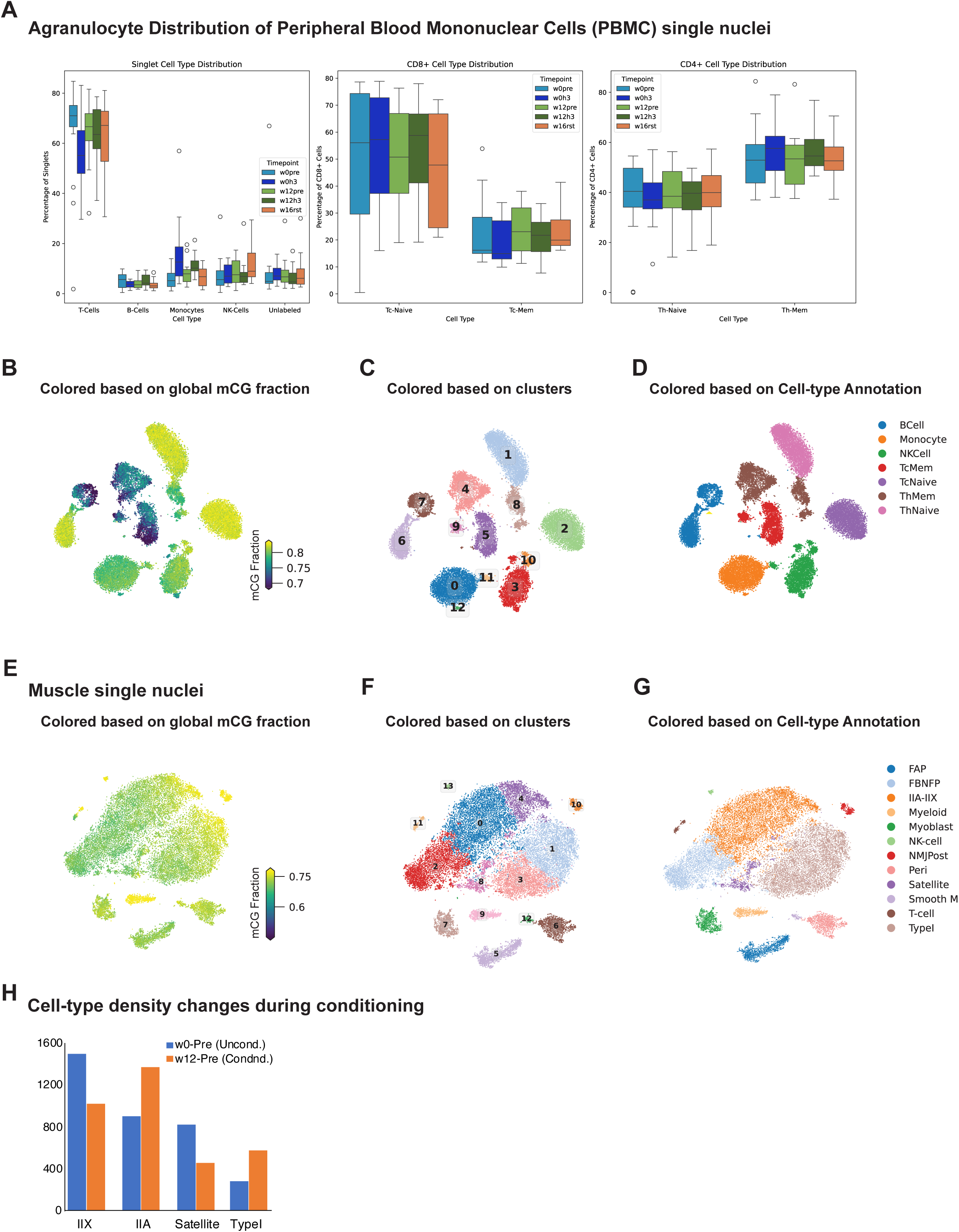
Single-nucleus Datasets. (A) The number of cells obtained from the various samples are grouped based on the timepoints and shown in the plots. The distribution of T-cells are much higher at all timepoints. They are then separated into Naive cells and Memory cells to see the differences of sub-populations of T-cells. (B) PBMC agranulocytes are separated into different clusters based on the global methylation levels at CG sites (C) and labeled as clusters. These can be further identified based on the antibodies used for FACS sorting and colored based on their cellular identity (D). Similarly, muscle cells are clustered based on global methylation levels (E), and labeled as different clusters (F). Using canonical marker genes, their cellular identities as assigned (G). In (H) the distribution of each of the muscle cell types are shown to compare their numbers at w0pre and w12pre timepoints. The distinct change in numbers have functional relevance.

DMRs were identified in each group of cell-types. Supplementary Table S5 summarizes the number of DMRs in each comparison. T-cells show maximum number of DMRs, irrespective of acute timepoints or conditioning response. Tc-Memory cells and Th-Naive cells show maximum response DMRs.

DNAm assay performed at single-nucleus resolution in muscle nuclei, provided distinct clusters of cells solely based on global methylation fraction across cytosine residues in the genome (Fig 7E, 7F, Supp Table S7). These distinct clusters could be annotated based on the hypomethylation patterns in the gene bodies of known marker genes for various types of myofibers such as MYH1 for Type IIX fiber, MYH4 for Type-IIB, PAX7 for satellite cells, and other cell types such as endothelial cells, immune cells, etc (Fig 7G). We were able to find a decrease of Type IIX fibers and an increase of IIA and Type I fiber after the 12-week training conditioning phase at the pre-exercise time point (Fig 7H).

We also identified exercise and conditioning responsive methylation changes (DMRs) within each cell type that offered more insights into the functional and developmental implications of the dynamic methylation patterns. We find that there are almost equal number of DMRs even at the acute response and detraining timepoints. This is probably due to the fact that we are specifically looking at various celltypes independently and not as a combined average of the whole tissue (Supp. Table S8).

We also identified differentially methylated genes (DMGs) based on gene-body methylation fraction. We found 481 DMGs in the Type IIA-IIX myofiber nuclei, 501 DMGs in Type I nuclei, and 223 DMGs in FAP nuclei. Since the numbers of DMGs are relatively low, we were not able to obtain function enrichment with adjusted-p-values < 0.05. However, with high odd-ratio and p-values < 0.05 we found that DMGs in Type IIAIIX are enriched in Vitamin D inflammatory diseases, pathways affecting IGF1 Akt Signaling, heart development, Gastric Cancer Network, Initiation Of Transcription And Translation Elongation HIV1 LTR as top five terms. DMGs in Type I were enriched in Development Of Ureteric Derived Collecting System, Alanine And Aspartate Metabolism. Neuroinflammation, Cardiomyocyte Signaling Converging On Titin. DMGs in FAP were enriched in Fatty Acids And Lipoproteins Transport In Hepatocytes, Familial Hyperlipidemia Type 5, Familial Hyperlipidemia Type 2, Type 1, and Primary Ovarian Insufficiency (Supp Table S9).

We noticed that the proportion of mature myofibers as well as progenitor cells (satellite cells) altered during the course of the 12-week exercise training, measured at baseline, pre-exercise timepoint. This effect has been reported in other studies ^100^. The number of IIX cells decrease as a result of the conditioning, whereas the number of Type IIA cells and Type I cells increase as a result of the consistent exercise. The proportion of stem cells decline considerably after the 12-week conditioning. This is probably due to the quick maturation and loss of stem-ness in the originally PAX7+, MYF5+ cells labeled as satellite cells.

## Discussion

Based on a sex-stratified cohort undergoing highly supervised, laboratory-based TRAD or HITT training we present the analysis of DNA methylation changes across 16 weeks, include two acute response studies completed at w0 and w12. Using blood and muscle, either as bulk tissue or at single nuclei resolution, we generated sequencing data at unprecedented levels (∼110,813,124 reads on average per sample; ∼88X coverage depth for bulk tissue, ∼2,613,755 reads average per nuclei in 26,187 PBMC, and ∼2,545,375 reads average per nuclei in 30,739 muscle nuclei). This enabled us to obtain base-resolution data required to quantify methylation status of individual cytosine residues.

The three comparison themes were chosen to match the study design and goal in assessing acute effects, chronic training adaptations, and sustained molecular changes that persist after stopping exercise training. The analysis presented here has shown distinct changes to the epigenetic landscape in blood as compared to muscle. While changes are much more pronounced in blood at the acute time points, the conditioning response is observed in muscle. These relate to the genomic regions where genes that respond immediately (inflammation, apoptosis, among others) and regions that respond to cell differentiation, stem cell maturation etc., in blood and muscle respectively. The epigenetic memory-associated DMRs are more prevalent in muscle. The timecourse data also offered the opportunity to study trends at baseline timepoints as seen by distinct patterns of methylation changes that are conserved across thousands of sites in the genome. The transcription factor motif enrichment also points to distinct pathways that would be regulated in response to acute exercise and chronic training adaptation. It is especially noteworthy that a distinct category of TFs are enriched in DMRs that persist even after a four-week period of detraining. These include TFs involved in satellite cell activation, myogenic differentiation and inflammation signaling (LEF1, PU.1, RUNX, EGR2, IRF8, SPIB, ETS). The correlation to phenotypic markers, although not universal, show strong correlations to more pronounced changes.

Single-nucleus data allowed us to delineate the various cell-types within the tissues. Cell-type-specific responses were prominent in PBMCs, especially in T-cells. The proportion of T-cell populations also were strikingly different from other agranulocyte populations. The agranulocytes were chosen in the study since they have the largest life-span and are the most responsive to stimuli. Muscle nuclei also revealed a heterogeneous population of various cell-type including the major myofibers such as Type IIA, IIX, Type IIB, Type I; endothelial cells, satellite cells, Fibro-adipogenic progenitors (FAPs) etc. Several distinct regions also were seen across these cell-types.

This study has a few key considerations. Due to sample availability for single nuclei sequencing, we were only able to test a subset of samples. We also note that while diet was not controlled across the complete trial, all baseline resting biospecimens were collected after an overnight fast, and all h3 tissue collections were matched using a post-exercise protein drink. The differentially methylated gene analysis depends on methylation levels averaged over the gene body. This measurement might not necessarily correlate to gene expression changes. We also point out that the function enrichment and motif enrichments point to specific pathways and biological processes, but not necessarily explain how the methylation changes will impact these functions.

Several studies have provided evidence that various factors contribute to the effectiveness of specific forms of exercise. Across ranges in age, sex, body type, and physical fitness, the intensity of training has heterogeneous effects on outcomes. Given the inter individual heterogeneity of response to exercise, it is challenging to recommend specific combinations of physical activity duration, intensity and frequency that would be optimal for a person given their goals, age, sex, comorbidities and behavioral patterns. Our study provides comprehensive insights into the DNA methylation changes associated with different exercise training regimens, training intensities, durations, and detraining periods. It highlights the complexity of epigenetic responses to exercise and sheds light on potential mechanisms underlying the health benefits of different exercise approaches. Building up on this and other large-cohort studies will invariably lead to actionable paradigms that can lead to better health outcomes, sustained throughout the entire lifespan of humans.

## Acknowledgements

This work was funded by the U. S. Department of Defense through the Office of Naval Research (ONR) (Fund number: N000141613159). We would like to thank all the unidentified donors who contributed biological samples for this project through our collaborators. The clinical trial site is at UAB Center for Exercise Medicine at University of Alabama at Birmingham (ClinicalTrials.gov Identifier: NCT03380923)

J.R.E. is an Investigator of the Howard Hughes Medical Institute. The shared resources in the lab is supported by grants from National Human Genome Research Institute (NHGRI) of National Institutes of Health (NIH) (Grant number: HG010634) Computational Infrastructure of Anvil HPC cluster at Purdue University through allocation MCB130189 from the Advanced Cyberinfrastructure Coordination Ecosystem: Services & Support (ACCESS) program, which is supported by National Science Foundation grants #2138259, #2138286, #2138307, #2137603, and #2138296, is acknowledged ^101^.

## Data Availability

The raw data and processed files are being submitted to a public repository.

The source code for various analysis is available at: https://github.com/manoj-hariharan/PHITE A web-browser with methylation data is accessible through http://neomorph.salk.edu/PHITE/index.php

## Methods

### Sample Donors

The whole-blood and muscle tissues were obtained from participants enrolled in the PHITE trial (Department of Defense, Office of Naval Research Grant: N000141613159, ClinicalTrials.gov Identifier: NCT03380923). The donors were of a similar age-group between 18–27 years and assessed as generally healthy, by questionnaires detailing health history and medical history, including screening for mental health disorders.

These participants were exercise-naive, as reported by no history of regular exercise training in the past 12 mo. All participants provided written, informed consent to participate and allowed their biospecimens to be used for future research. More details can be obtained from the collection center at The University of Alabama at Birmingham (UAB).

The participants were randomized and assigned to either traditional combined exercise training (TRAD group with 27 participants - 15 females and 12 males) or high-intensity tactical training (HITT group with 27 participants - 16 females and 11 males). Randomization was stratified by sex to ensure similar distributions of females and males in TRAD and HITT.

Metadata associated with the donors are provided as Supplementary Table S1.

### Bulk-tissue Methylation Profiling

DNA was extracted from whole blood samples (100ul), or muscle samples (∼10mg), using DNeasy Blood and Tissue kit from Qiagen and resuspended in 125ul of EB buffer.

The Methy-C-seq libraries were constructed following the SureSelect Human Methyl- Seq™ Target Enrichment System for Illumina® Multiplexed Sequencing protocol from Agilent Technologies® for low input DNA, the protocol for samples for 1ug of sample, instead we used 750ng of DNA to start the libraries, we follow the protocol from the fragmentation to the capture, we also used the EZ DNA Methylation- Gold Kit from Zymo for the bisulfite conversion, and we did 2 sets of PCR amplifications following the protocol set up, one to add the common primers and the second one to add the index primers for pooling and sequencing (9+6 PCR cycles). The libraries were quantified using Qbit by Invitrogen® and TapeStation and High Sensitivity D1000 ScreenTape by Agilent Technologies®. MethylC-seq libraries were sequenced on Illumina® HiSeq 4000.

These are based on capture probes designed specifically to focus on 91% of all CpG islands, cancer- and tissue-specific DMRs reported in previous studies, promoter regions of genes, gene bodies, and other regulatory regions defined in the ENCODE project.

Quality Control metrics for the assay are provided in Supplementary Table S2.

### Bulk-tissue data analysis

Raw methylation data were obtained as individual allC files for each sample. Each patient provided matched muscle and blood tissue samples across five distinct timepoints, resulting in separate allC files per patient, per tissue, per time point. Individual allC files were first merged within groups defined by Sex, Tissue, Intensity, and Time Point using the merge-allC command from the ALLCools package ^36^. This merging step aggregates methylation counts across samples within each subgroup into a single allC file representing the subgroup for use in the next step of identifying differentially methylated regions (DMRs).

DMRs were identified using the methylpy software package ^37^, which applies a statistical approach to detect genomic regions showing significant methylation differences between conditions. Specifically, pairwise comparisons were performed between timepoints within subgroups of distinct combinations of sex, tissue, and training intensity. Of the five unique timepoints, we crafted six individual comparisons representing exercise response, conditioning response, and long-term response. We defined exercise response as the comparison between timepoints before training (w0pre or w12pre) and three hours after training (w0h3 or w12h3). We defined conditioning response as the comparison between timepoints before (w0pre or w0h3) and after (w12pre or w12h3) the course of training. We defined long-term response as the comparison between baseline timepoints just before (w0pre) and after the course of training (w12pre) with the time point after stopping the training regiment (w16rst). Thus, our pairwise comparisons were as follows: w0pre vs w0h3 and w12pre vs w12h3 for exercise response, w0pre vs w12pre and w0h3 vs w12h3 for conditioning response, and w0pre vs w16rst and w12pre vs w16rst for long-term response. For each pairwise comparison, allC files from each subgroup and time point were input into methylpy DMRfind.

Prior to statistical testing, methylpy filtered cytosines to include only positions with sequencing coverage of at least 10 reads (min-cov=10) across both comparison groups. Analysis was restricted to cytosines within the CG context (mc-type="CGN"), thereby focusing exclusively on CpG dinucleotide sites. Methylpy then applied a permutation-based root-mean-square (RMS) statistical test (run_rms_tests), comparing methylation counts at individual cytosine positions between the two groups. Specifically, observed methylation differences were assessed against a distribution generated from 3,000 permutation simulations (num_sims=3000). FDR was estimated from the p-values generated by the RMS test based on a histogram approach. This iterative algorithm estimated the number of true null hypotheses and determined an adjusted p-value threshold corresponding to an FDR cutoff of 1% (sig_cutoff=0.01). Cytosines that passed this test, also defined as differentially methylated sites (DMS), were subsequently merged into contiguous genomic blocks, or DMRs, by grouping significant cytosines within 250 base pairs of each other (dmr_max_dist=250). Regions were required to contain consistent directional changes in methylation (i.e., hyper- or hypo-methylation) across samples within each group. This was enforced by evaluating the magnitude and direction of residuals from expected methylation levels at each cytosine position. From the residual distribution, sites in the right tail were considered hyper-methylated, while sites in the left tail were considered hypo-methylated. The resulting DMR output contained genomic coordinates, the number of DMS in each DMR, an indication of which group in the comparison the DMR was hyper- and hypo- methylated, as well as the fractional value of the mC signal divided by coverage of the DMR for each group.

For each DMR file, we appended the mC fraction values of each sample in both subgroups for each DMR. To do this, we utilized allCools._allc_to_region_count.batch_allc_to_region_count to generate allC files specific to the DMR regions, containing the sum of the CGN context mC counts and sum of the coverage counts across each DMR. We then map each DMR from the allC to the DMR file produced by methylpy DMRfind in order to add the fractional methylation values of each sample.

We conducted a two-tailed independent t-test for each DMR to evaluate whether methylation fractions differed significantly between the two conditions being compared. Prior to statistical testing, we applied a stringent missing data filter to enhance reliability. Any DMR that failed to retain at least 40% of its original samples in either condition (i.e., at least 60% missing methylation values) was filtered out.

For each DMR that passed this threshold, we calculated a raw p-value using an independent two-sample t-test comparing fractional methylation values between samples of the two groups. To control the false discovery rate due to multiple comparisons, we applied the Benjamini-Hochberg (BH) correction to generate adjusted q-values. DMRs passing stringent significance criteria (raw p-value ≤ 0.05 and adjusted q-value ≤ 0.1) were retained as statistically validated DMRs. DMRs exhibiting less than a 10% absolute difference in fractional methylation between the two comparison groups were also removed to retain only significant methylation changes.

For identifying enriched transcription factor binding site (TFBS) motif enrichment, we used HOMER within 50nt around a DMR if the length of the DMR is less than 100nt or the entire length of the DMR. Here we present only the motifs that are enriched above a FDR corrected q-value < 0.05 with 440 known human TF motifs.

For DMG analysis, the first step is to quantify methylation at the gene level. We defined genomic regions encompassing each gene, extending 2 kb upstream and downstream (±2 kb) of transcription start and end sites, respectively. These expanded genomic coordinates were generated from the GENCODE v19 reference annotation for the human genome (hg19).

We then calculated methylation fractions for each gene-region across individual samples by utilizing the batch_allc_to_region_count function from the ALLCools package. Methylated cytosine counts and total cytosine coverage counts within the CGN context were aggregated across each defined gene-region for every sample. Fractional methylation was calculated as the ratio of aggregated mC counts to total cytosine coverage for each gene-region and appended to create gene-level methylation profiles.

Using the gene-level methylation profiles generated above, we identified genes exhibiting significant methylation differences between conditions, referred to as differentially methylated genes (DMGs). Prior to differential testing, methylation fraction matrices were filtered to retain only protein coding genes with at least 90% non-NA sample methylation fractional values. Remaining missing methylation fraction values were imputed using distance-weighted K-nearest neighbors (KNN) imputation (n_neighbors=5, weights="distance"). Following imputation, the methylation data were scaled, and principal component analysis (PCA) was performed to reduce data dimensionality. Differential methylation analysis was conducted by comparing methylation fractions between conditions at each gene using Scanpy’s rank gene groups functionality (sc.tl.rank_genes_groups) ^103^. Specifically, we employed a t-test with overestimated variance (method=’t-test_overestim_var’) to robustly identify significant methylation differences while controlling false positives. BH correction was used to generate adjusted p-values and genes with adjusted p-values below 0.05 were considered significantly differentially methylated.

### Longitudinal Variation

B-splines are a statistical modeling technique to fit a polynomial curve using a basis function between points. Because we had several replicates per time point, the B-spline approach with a basis function was deemed most appropriate to detect significantly changing splines. To then compute significance, a null model of a flat change in methylation proportion was centered around the B-spline’s intercept, and a residual squares test was used to compute a p-value between the null model and the spline’s model.

To cluster together the tens of thousands of splines, we used K-Means clustering with a maximum of 300 iterations, and a minimum of 10.

To compare the cluster specificity of certain regions, we employed a normalization scheme where after binning the counts per genomic region, we normalized these by each cluster’s size and set a scaling factor of 10,000. For computing correlations between normalized counts sites, a permutation test was used with n=10,000, where the counts were randomly shuffled in the bins. Computing cluster-specific sites was performed by multiplying two metrics: the inverse entropy of each binned region using the normalized counts, and the difference between the maximum and the next highest score. Using both an entropy metric and a metric based on the absolute value of counts allows us to find both distinctive sizes from the entropy, and large sites with the latter metric. We then performed a z-score normalization on this multiplicative score, which is the final cluster-specific score.

### Single-nucleus Methylation Profiling

Sequencing libraries were prepared using the protocol called Single-nucleus methylation sequencing v2 (snmC-seq2) described in [^104^ and ^101^]. Briefly, the DNA undergoes bisulfite treatment for C-T conversion and barcoded with random primers. The samples are then pooled through two rounds of SPRI cleanups such that 16 384-well plates are compressed to a single 96-well plate. The pooled samples are then amplified and further purified with two more rounds of SPRI cleanups. After confirming library concentrations and normalized, these libraries will proceed to sequencing. Sequencing was performed on S4 flowcells in the 150-bp paired-end mode on Illumina™ Novaseq 6000™.

### snmC Sequencing Data Processing

snmC-seq data processing is done by following a custom pipeline called YAP v1.6.9 as previously described [^105^] [https://github.com/lhqing/cemba_data]. The main steps include (1) demultiplexing the FASTQ files for the respective single nuclei based on the primer information of each well of the 384-well plate using cutadapt v4.4, (2) read-level quality control (QC) using samtools v1.20 and picard v3.0.0, (3) mapping to the mouse hg19 genome using bismark v0.24.2 and bowtie v2.5.4, (4) processing mapped BAM file to generate QC metrics, and (5) generate base-level molecular data of methylation profiles as outlined in the “Dataset Organization” section below, using allcools v1.0.23.

### Quality Control Metrics

We use three metrics to filter out nuclei: (1) mCCC fraction: this is a proxy measure for the upper bound of the C-T non-conversion rate. It is the methylation fraction of C in the CCC context. This value closely mimics the C-T non-conversion rates, as measured with lambda DNA spike-in, which is unmethylated. We filter out nuclei that exceed a 5% threshold. (2) Mapping rate: the Bismark mapping rate value gives an idea of possible contamination or wells that did not have any nuclei loaded. We filter out nuclei with less than 50% mapping rate. (3) total final reads after removing any duplicates should be between 500,000 and 10,000,000 for any nucleus used.

### Dataset Organization

The single-nucleus methylome profiles obtained after mapping are stored in a simple tab-delimited file called All Cytosine (allC) format which is indexed by bgzip/tabix ^106^. It is a six-column format containing the methylation information for every cytosine residue that has been sampled. It contains the chromosome number, coordinate, strand, three-base sequence context, count of reads supporting methylation, and total number of reads (coverage).

We also generate a cell-by-feature matrix of methylation profiles using the methylated base counts (mC) and total counts (cov) for each position or a region. This data matrix is stored in a modified xarray dataset file called Methylcytosine dataset (MCDS) file. It stores four common dimensions representing cells, regions, context of the cytosine, count type (mc or cov).

Details of the allC and MCDS format and organization are provided in allCools (https://lhqing.github.io/allCools/start/input_files.html).

### DNA methylation-based Clustering

The methylation profiles obtained by aggregating the mC and cov values in non-overlapping windows of genomic bins of 5kb for each nuclei are defined as a chrom5k region. We generate separate chrom5k MCDS dimensions for methylated cytosines in the CG and CH (C[ATC]) contexts. This approach aggregates single-nucleus profiles of large genomic regions thereby allowing increased coverage while eliminating the need to access several terabytes of single-nucleus level files at various analysis stages. These were used as input for identifying clusters of nuclei which have common methylation profiles within them and separate across the clusters. We performed methylation clustering iteratively using the pipeline described in ^106^.

Briefly, we perform the following six steps using the chrom5k matrices: (1) removing bins that overlap ENCODE blacklist [^101^], (2) selecting highly variable features (HVFs), n=5000, from CG-methylation using support vector regression (SVR), (3) generating posterior mCG fraction matrices for each bin described in ^108^, (4) clustering with HVF and calculating Cluster Enriched Features (CEF), adapted from the cytograph package ^109^, (5) calculating principal components (PC) in the selected cell-by-CEF matrices and generating the t-SNE ^110^ and UMAP ^111^ embeddings for visualization. t-SNE was performed using the openTSNE package ^112^, (6) consensus clustering: we first performed Leiden clustering ^113^ 200 times using different random seeds and combined the labels to establish preliminary cluster labels. We then trained a predictive model in the PC space to predict labels and compute a confusion matrix. This enabled us to merge similar clusters and minimize confusion, based on the SCCAF package as implemented in ^102^. This iterative process continued till the model’s performance on withheld test data achieved an accuracy of 0.95 for the initial round and > 0.9 for all subsequent rounds. This framework is implemented in the allCools module “clustering.ConsensusClustering”. These steps also include various packages from scanpy ^114^ and scikit-learn ^115^.

### Cell-type Annotations of Clusters

As previously described, we generate MCDS also using gene bodies as regions. The gene body region is defined as the region starting 2kb upstream of the TSS and ending after 2kb downstream the TSE. We use the gencode v19 reference annotation which corresponds to the hg19 genome.

An inverse relationship is known to exist between methylation level of the gene body and the gene’s expression. Although this is not a standard rule, and can vary across cell-types, cell-state, and other regulatory elements, we explored this possibility.

We examine the gene-body regions’ methylation for well-known (canonical) marker genes for the various myofibers, and other cell-types known to be associated with muscle dissections (such as adipocytes, endothelial cells, myeloid cells etc.).

Based on the unique and strong hypomethylation profiles of such canonical marker genes, we were able to assign a cell-type label for all clusters.

### Cell-type specific Merge of Nuclei

After assigning cell-type labels to individual nuclei from the clustering analysis, we merged individual single-cell allC files into the pseudo-bulk level by utilizing the merge- allc command from the allCools package. This command takes a list of individual allC files and, for each unique coordinate across all files, sums up the mC signal and coverage signal at each coordinate. We merged allC files based on matching sexes, intensity, timepoint, and cell types.

### Identifying Differential Methylation from Clusters of Single Nuclei

In order to identify sites or loci in the genome that had statistically significant differences in the methylation profiles comparing the timepoints in acute phase or condition we implemented the pipeline which is also implemented in allCools. The pipeline is referred to as Differentially Methylated Region (DMR) calling. Before DMR calling, we omitted pseudo-bulked clusters which have less than 30 nuclei in either of the conditions. We then performed DMR calling using the allCools.dmr.call_dms to identify Differentially Methylated Sites (DMSs) and, subsequently, allCools.dmr.call_dmr to identify DMRs from the identified DMSs. Individual DMSs were first identified with call_dms, in which for each methylation site specified in the pseudo-bulked allC files, a Root Mean Square (RMS) test with permutations is performed to assess significance of methylation differences, and the p-values are stored for each site. A minimum p-value of 0.01 was applied to the identified DMSs. We decreased this p-value threshold to 0.001 while running call_dmr, in which we only consider DMSs of p-value less than 0.001 and a change in methylation fraction of 0.3, in which they are grouped into DMR windows with a maximum allowable distance of 250 base pairs. For multiple DMSs within a DMR window, a Pearson correlation of their methylation change across samples is calculated, and only DMSs that have correlation above a cutoff of 0.3 are grouped together. DMSs within the same window that do not meet the correlation threshold are split into separate DMRs. Each DMS is classified as hypo- or hyper-methylated based on the residuals of their methylation fractions. We set a residual quantile cutoff of 0.7, meaning that hyper-methylated DMSs have a residual quantile greater than 0.7, and hypo-methylated DMSs have a residual quantile less than 0.3. Finally, the counts and fractions (frac) of each DMR is recalculated and saved in Zarr format.

Once the DMR calling is complete, we load the Zarr formatted output into a RegionDS dataset. We filter the DMRs using bedtools’s intersectBed with a blacklist, and remove all DMRs within the blacklisted regions. We then split the RegionDS object into hypo- and hyper-methylated objects in relation to the Sed condition, and load them both into pandas. We then save the chromosome, start position, end position, number of DMSs, length of DMR, and sample DMR frac fields into a tsv.

### Merge-allC by replicate

Using the merge-allc command from the allCools package again, we then merged allC files based on matching sexes, intensity, timepoint, and cell types to obtain replicate specific allC files.

### Retrieving DMSs

To retrieve the exact DMS positions that make up each DMR, we reimplemented allCools.dmr.call_dmr function and had it save the individual DMS position information when concatenating DMSs to DMRs. To do this, we load the allCools.dmr.call_dms output into a RegionDS object. We perform the same filters described in the Call DMR step, however, we save the DMS Positions as a list in the object, as well as the DMR IDs. Finally, we create a new DMS object from each position in the DMS Positions field, saving chromosome, start position as 0-based, end position as 1-based, and the DMR ID. We saved this information into a tsv.

### Add mC level

For each DMS file, we added the mC fraction values of the replicates for each DMS. To do this, we utilized allCools._allc_to_region_count.batch_allc_to_region_count to generate allC files specific to the DMS regions, containing the sum of the CNN context mC counts and sum of the coverage counts across each DMS. We then map each DMS back to its DMR using the DMR ID field, and calculate the fractional value of each sample from the sum of each DMSs’ mC divided by the sum of each DMSs’ Coverage in each DMR.

### Predictive Model Development

The predictive models were developed based on Random Forest Classifier (RFC) which is an ensemble machine learning method. The three major aspects of this method is given below.

Feature Selection: We implemented an iterative approach to identify the most informative DMRs of each comparison across the training groups in the respective tissues, for 200 iterations. We used stratified 3-fold cross-validation (CV) in every round to train the RFC. The importance of each feature (DMR) relative to the condition and the donor are identified. The importance of the remaining features are gauged by the average feature importance derived from the RFC. The top 200 features were chosen in every round and the rest were reserved for the next round. The RFC uses 500 trees with a maximum depth of 3 for each iteration.

Mitigating Donor Effect: In order to mitigate the biological effect of donor variability due to confounding factors, we selected shared features from the prior 3-fold CV as candidates. We then evaluated their importance in predicting the either the given conditions of the comparison groups or in determining the donor. The scatter plot in figure 5A is a representative visualization of the cumulative feature importance across the iterations. We utilize only those features that held a positive importance for the comparison groups and an importance of less than 5e-4 for donor prediction. The density estimates towards the left within the yellow highlighted region are the features that would be selected.

Method Evaluation: For assessing the predictive capability of the selected features, we implemented 4-fold CV using the features stratified based on the combination of the conditions being compared. In each fold, we used three subsets of the features as feature selection process as outlined earlier. The RFC was trained based on the chosen feature which was then used on the remaining fold to determine the accuracy of the model. The model accuracy for each of the 108 comparisons are summarized in Fig 4B.

### Phenotype Correlations with DMRs

We visualized average methylation levels of differentially methylated regions (DMRs) for each defined group (combining gender, intensity, exercise, and tissue) using the PyComplexHeatmap ^115^. Within each group, we employed Wilcoxon rank-sum tests to identify phenotypes exhibiting significant differences in measurements between two comparison groups, as displayed in the corresponding heatmaps. Subsequently, for each DMR (both hyper- and hypomethylated), we calculated the Spearman correlation with the measurements of the significant phenotype across the relevant samples. The reported correlation coefficients represent the median of coefficients for DMRs with a Spearman p-value ≤ 0.05.

