## Supplementary figures and images for "DNA Methylation Dynamics of Dose-dependent Acute Exercise, Training Adaptation, and Detraining"

### Supp. Fig. S1

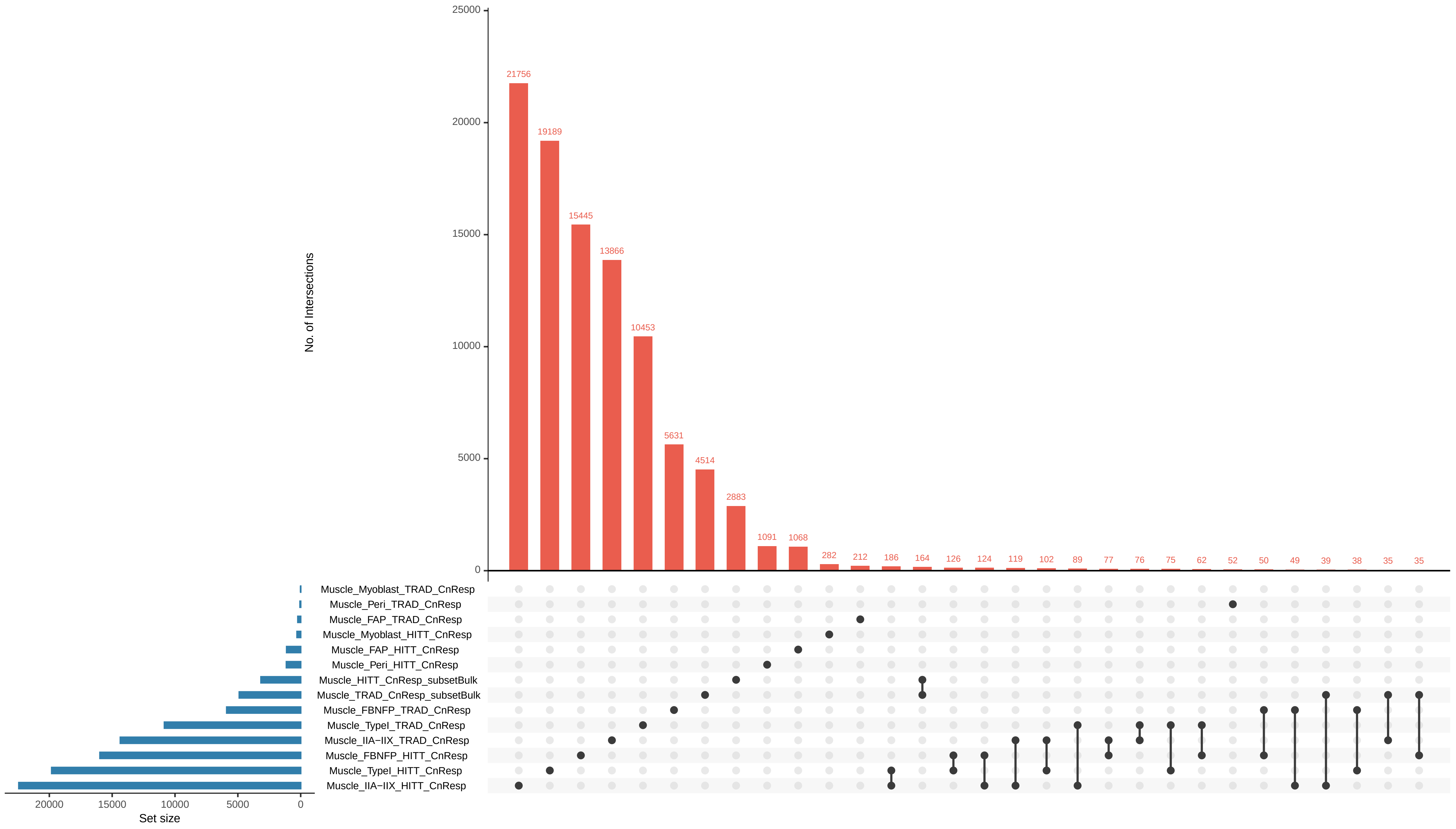

### Supp. Fig. S3

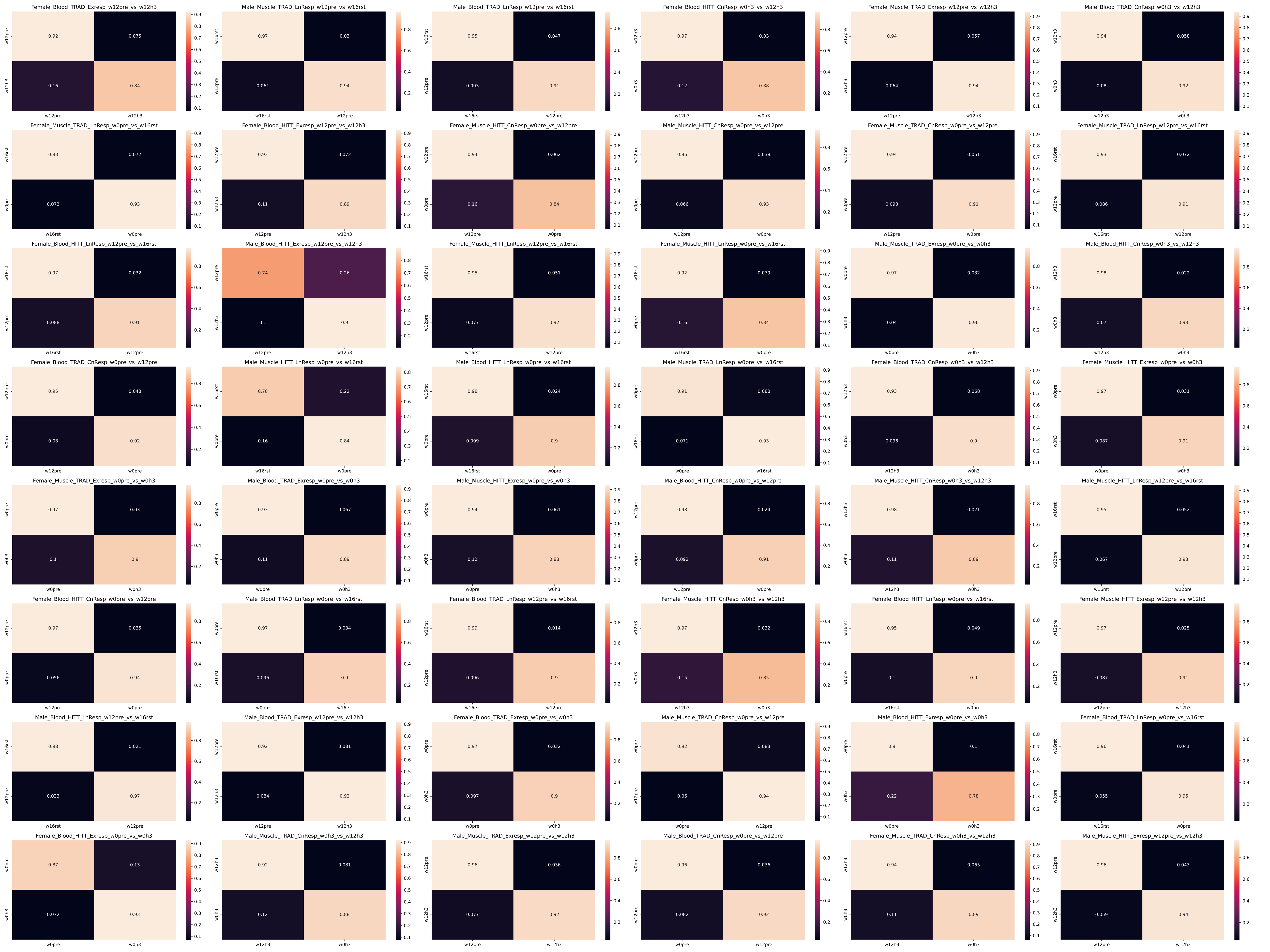

### Supp. Fig. S4

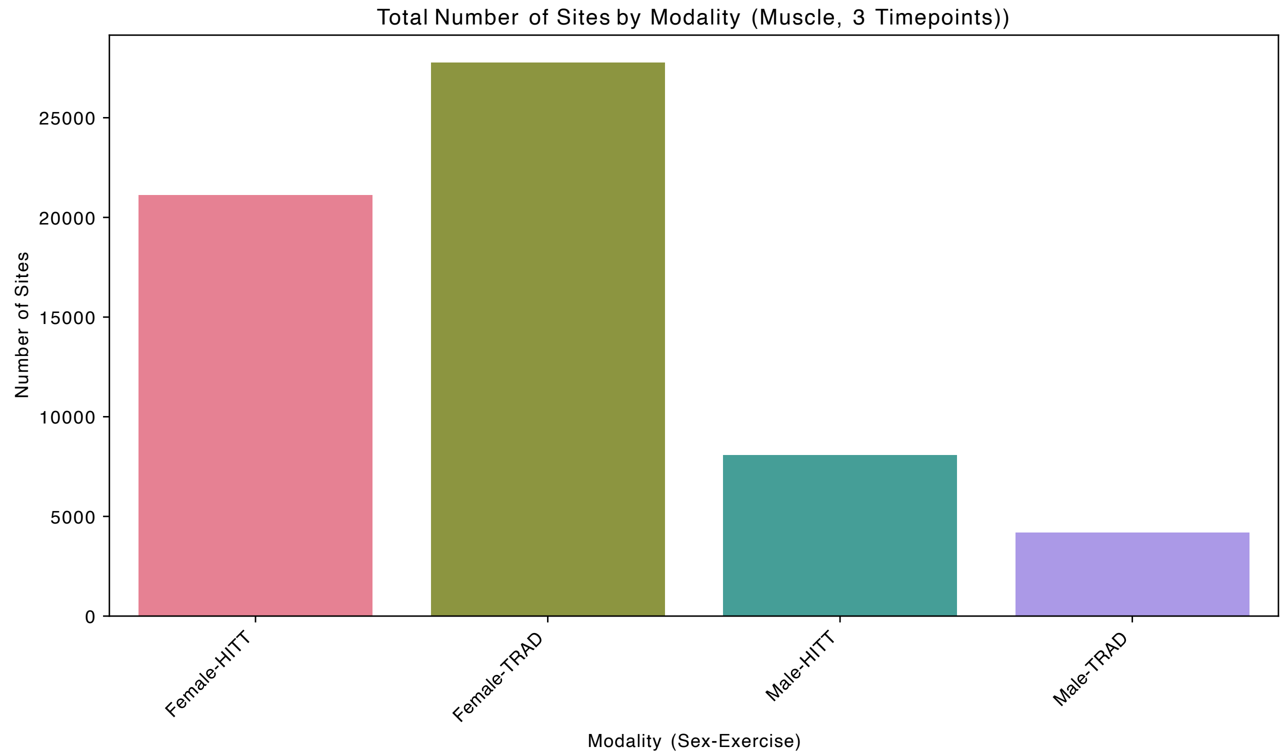

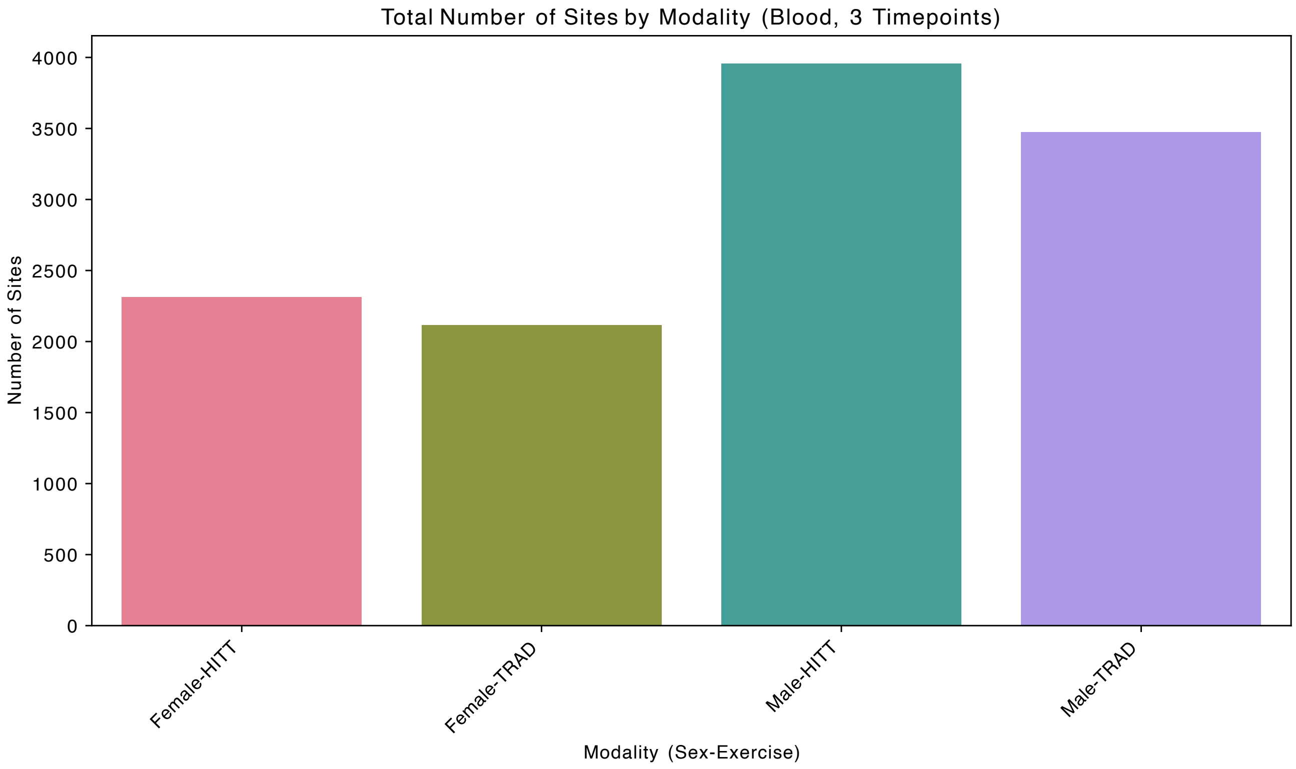

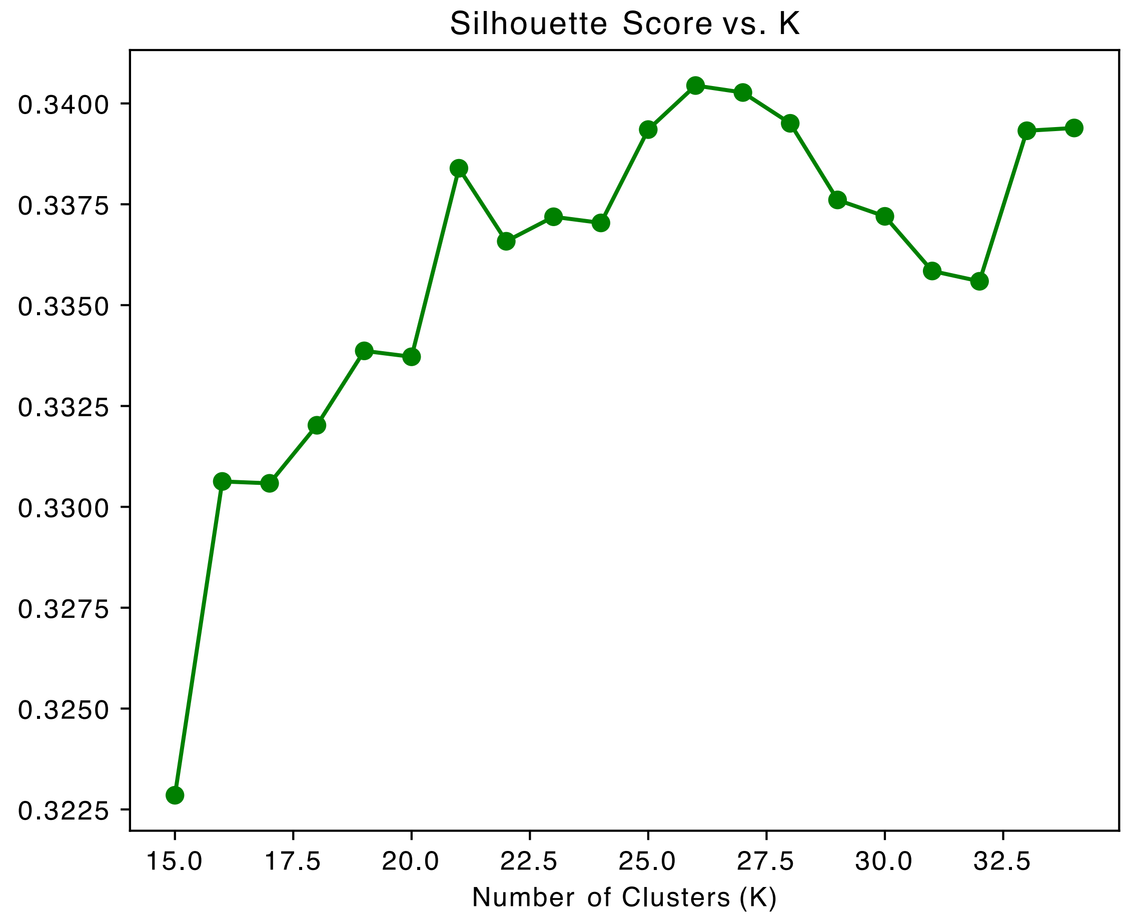

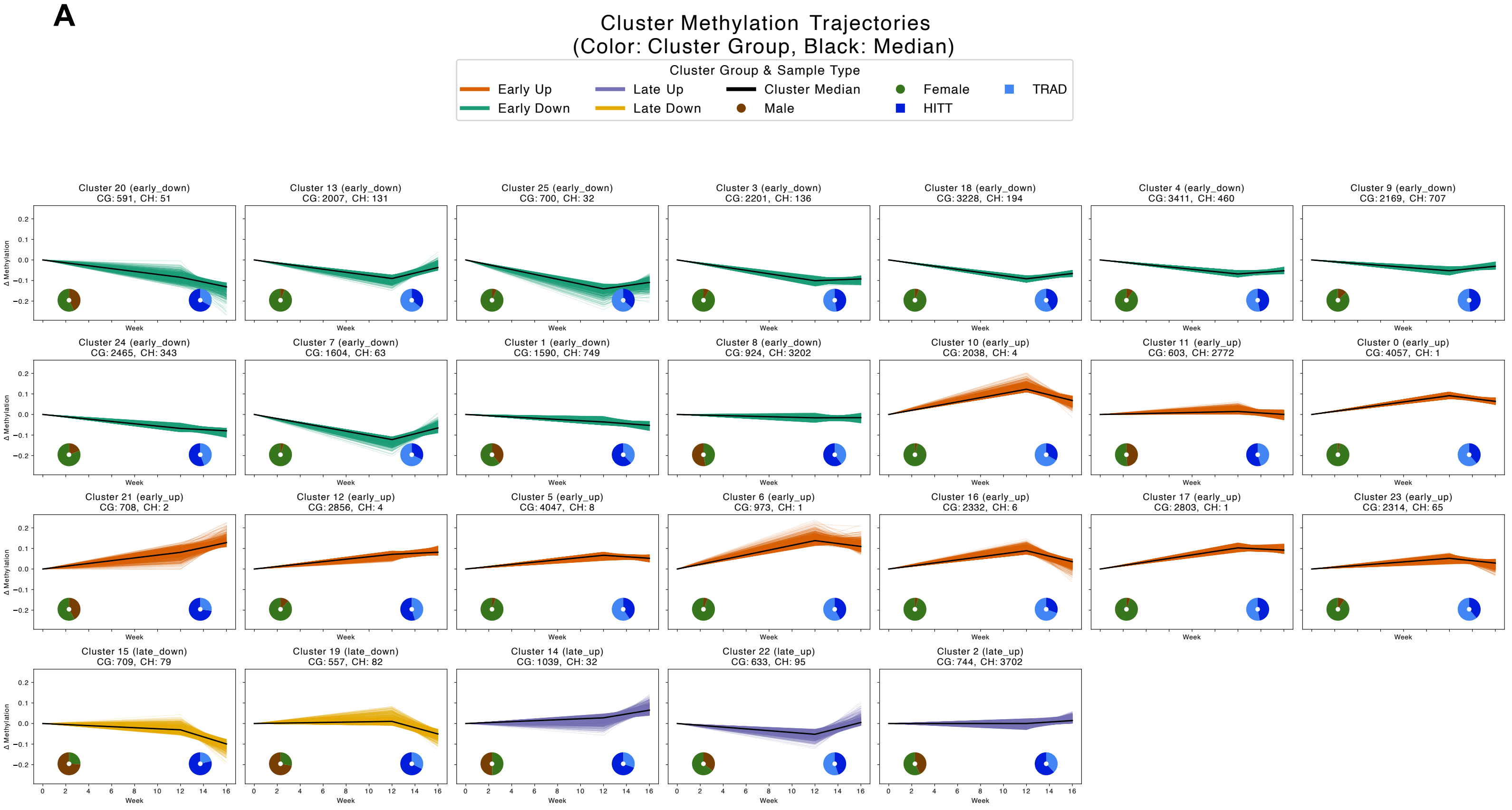

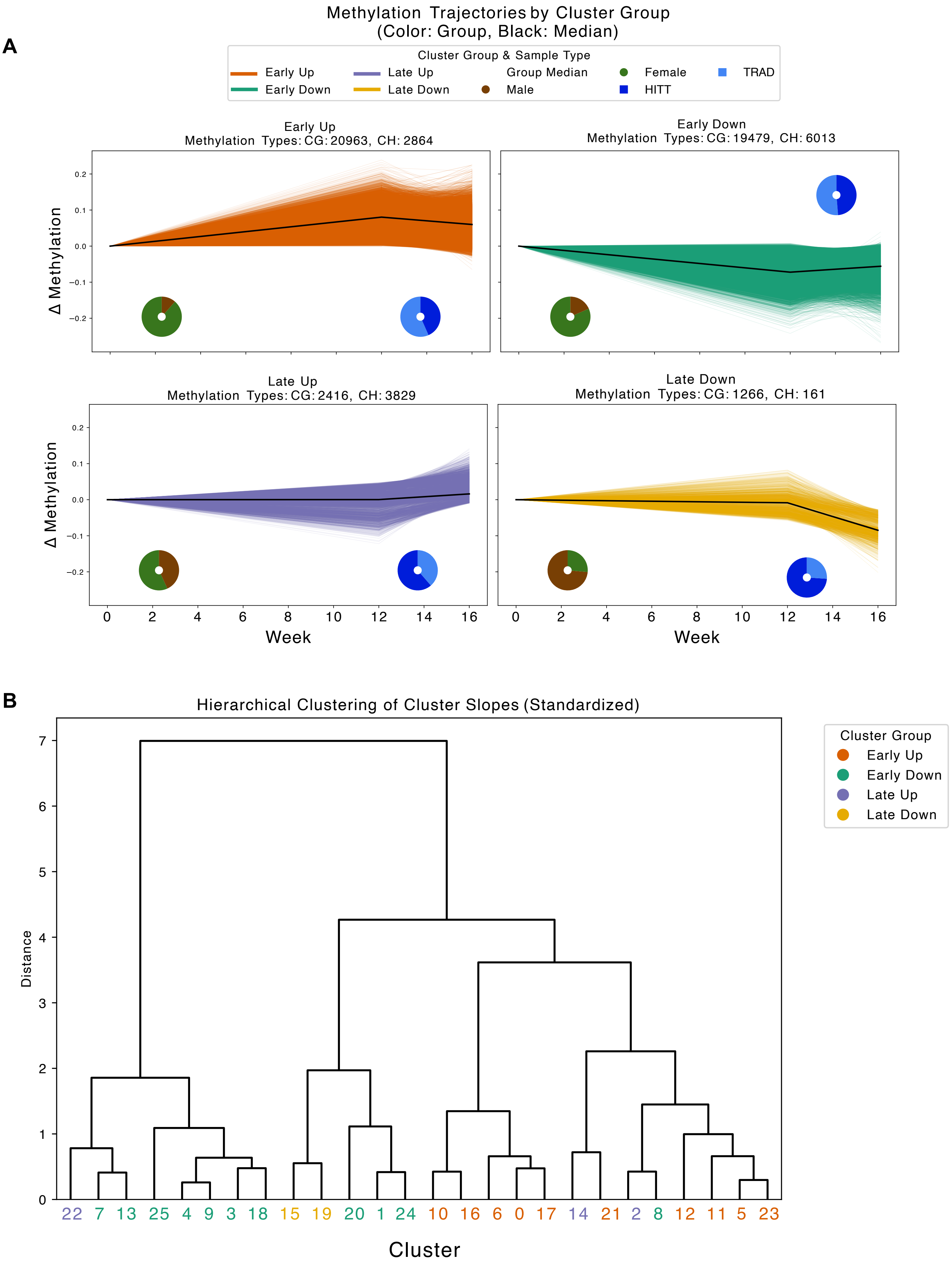

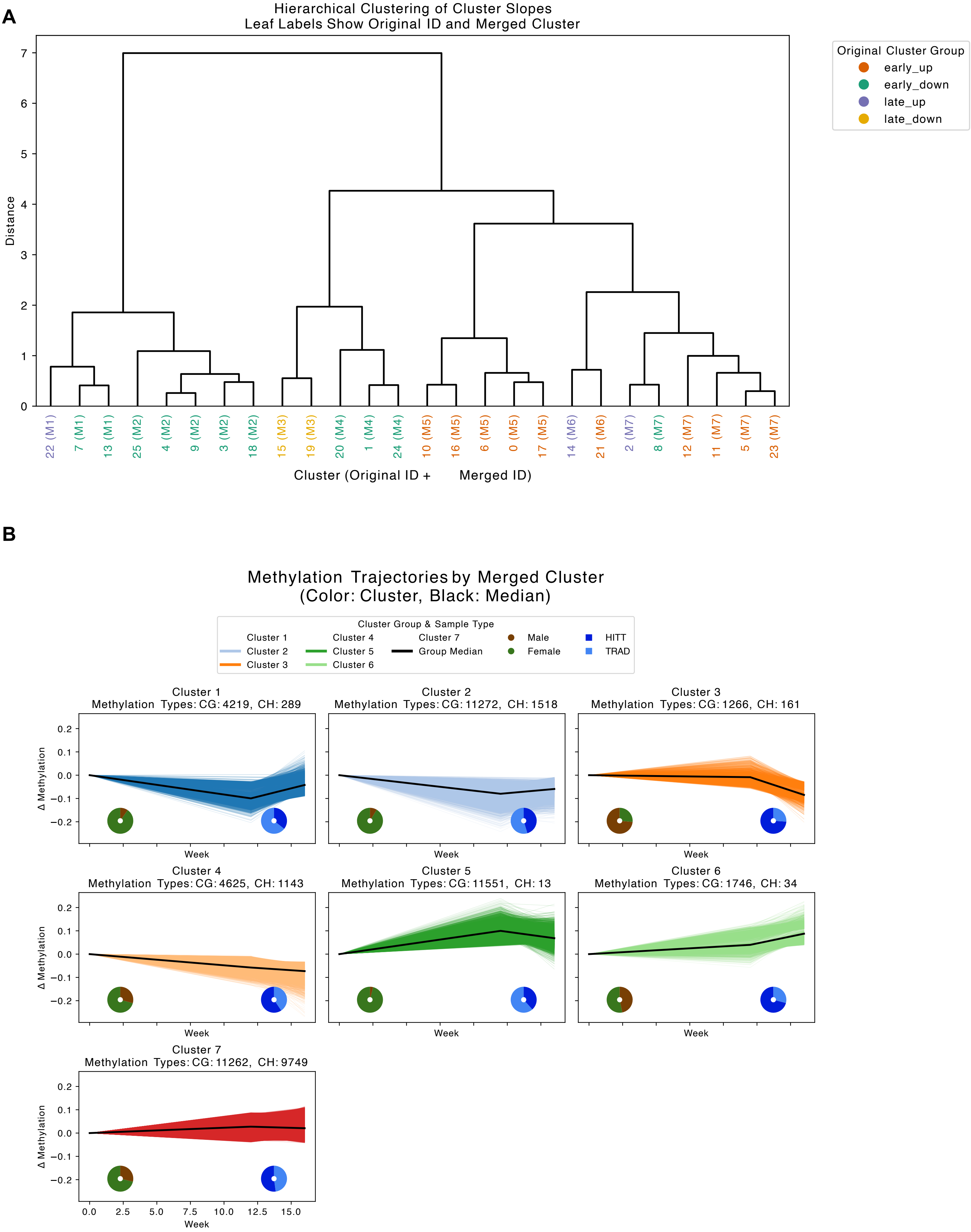

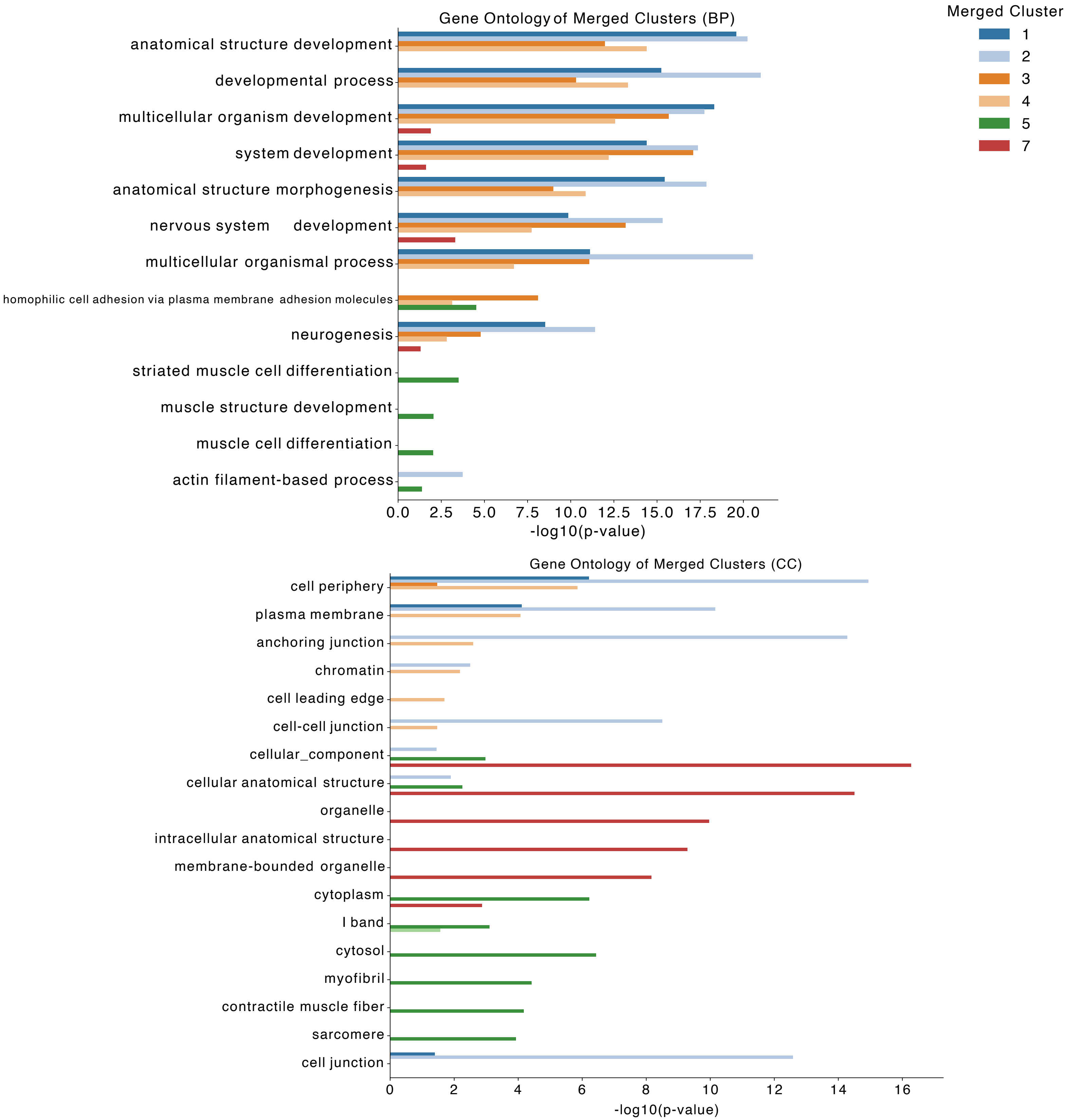

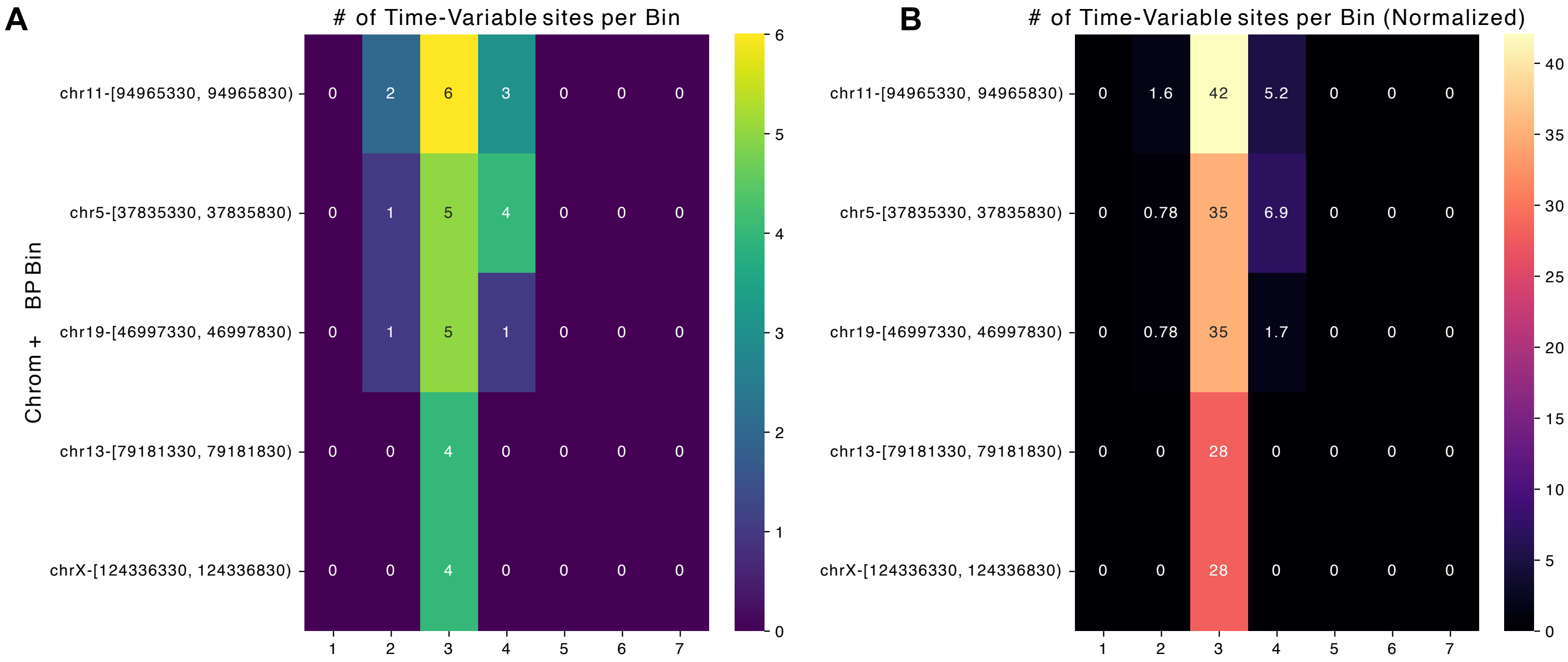

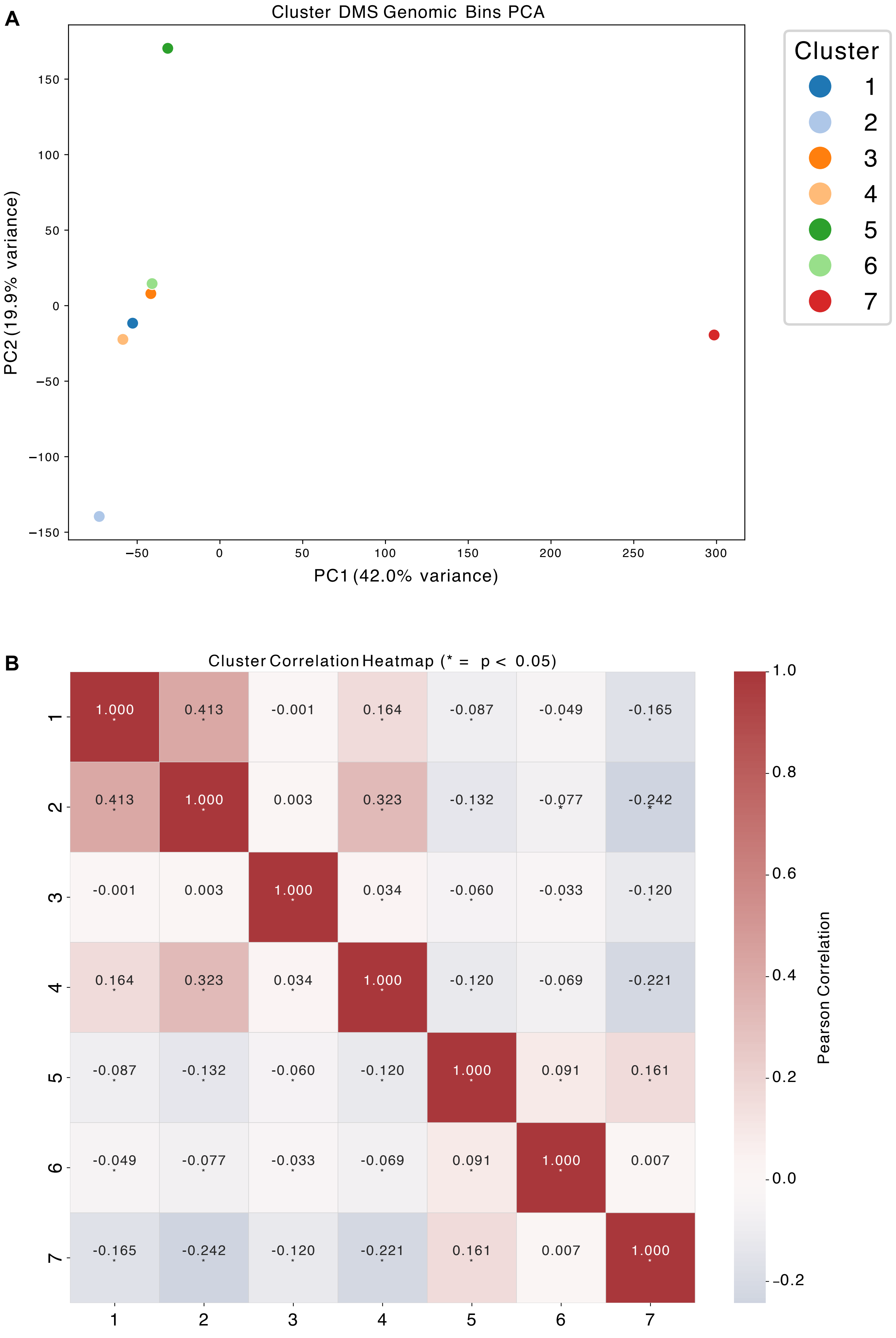

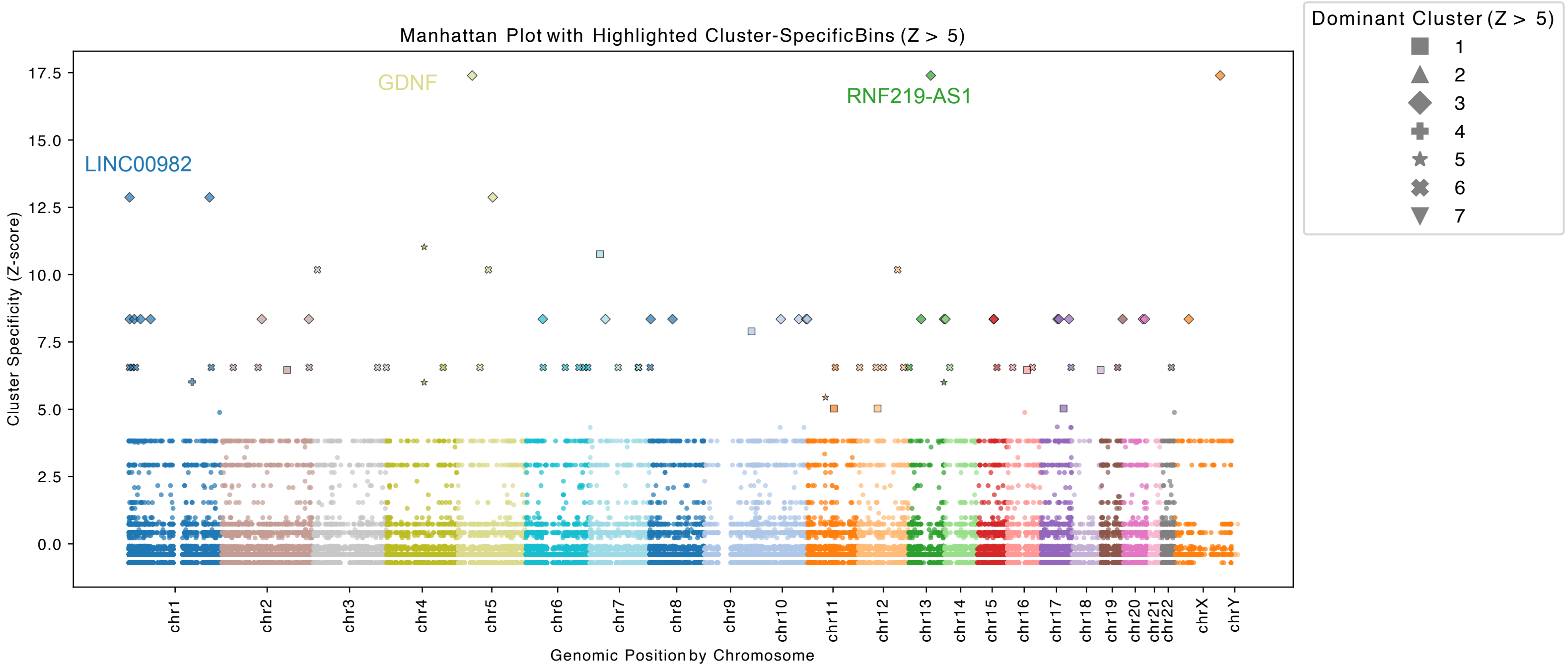

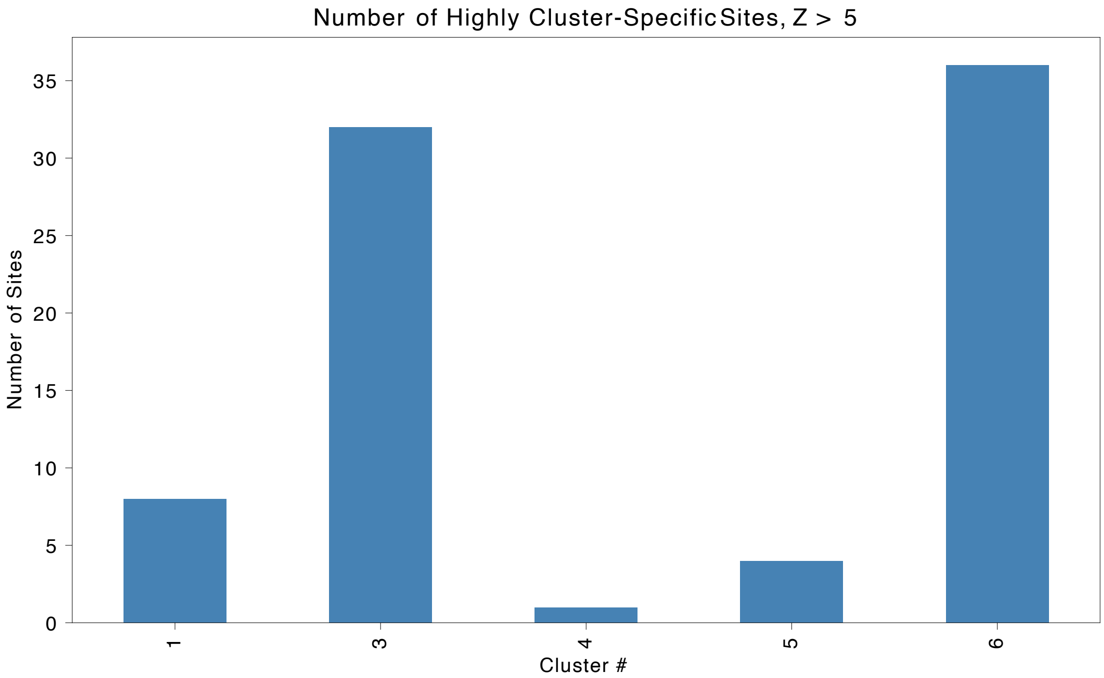
