## Supplementary material for "DNA Methylation Dynamics of Dose-dependent Acute Exercise, Training Adaptation, and Detraining": Supp. Fig. S2

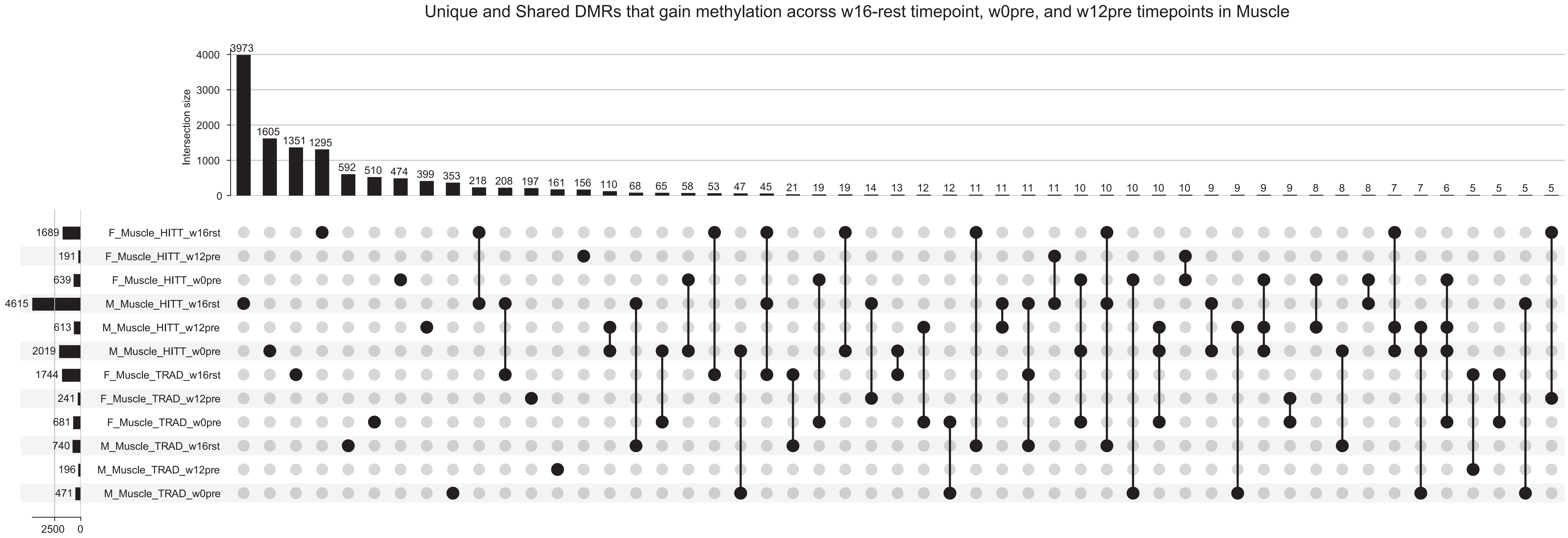
Unique and Shared DMRs that gain methylation acorss w16-rest timepoint, w0pre, and w12pre timepoints in Muscle
Intersection size
3973
1605
1351
1295
592
510
474
399
353
218
208
197
161
156
110
68
65
58
53
47
45
21
19
19
14
13
12
12
11
11
11
11
10
10
10
10
10
9
9
9
9
8
8
8
7
7
6
5
5
5
5
F_Muscle_HITT_w16rst
F_Muscle_HITT_w12pre
F_Muscle_HITT_w0pre
M_Muscle_HITT_w16rst
M_Muscle_HITT_w12pre
M_Muscle_HITT_w0pre
F_Muscle_TRAD_w16rst
F_Muscle_TRAD_w12pre
F_Muscle_TRAD_w0pre
M_Muscle_TRAD_w16rst
M_Muscle_TRAD_w12pre
M_Muscle_TRAD_w0pre
1689
191
639
4615
613
2019
1744
241
681
740
196
471
2500
0

### Unique and Shared DMRs that lose methylation acorss w16-rest timepoint, w0pre, and w12pre timepoints in Muscle

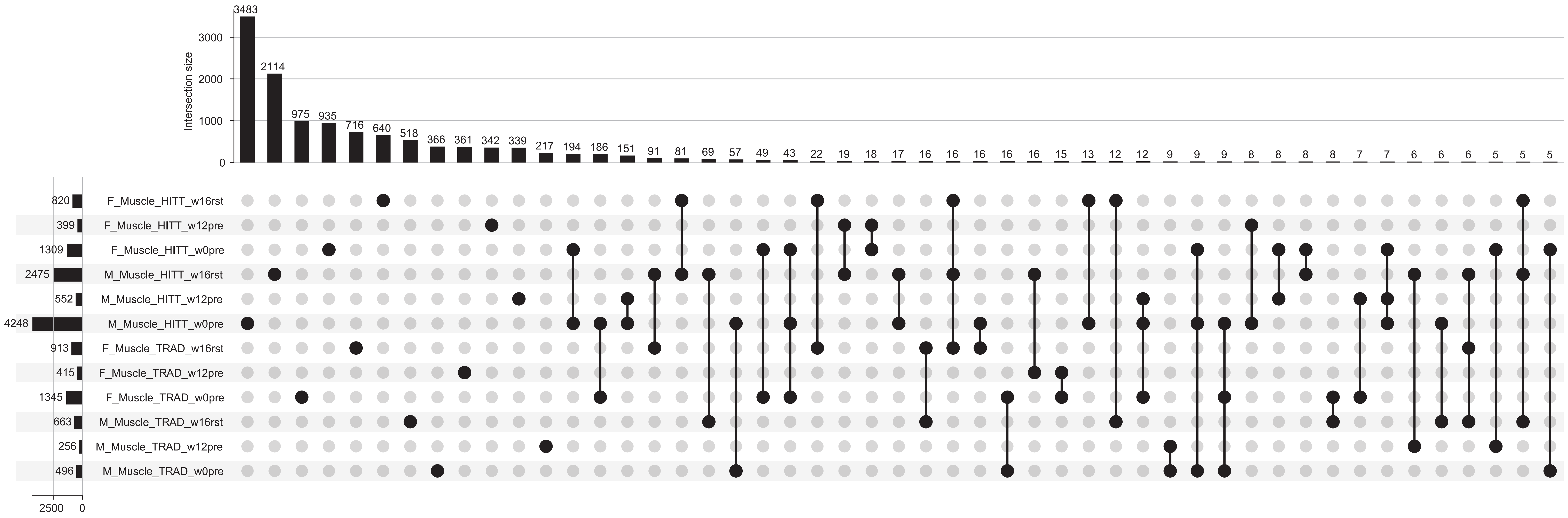
